# MorphoProt: RNA-Conditioned Integration of Multiscale Histology for Spatial Protein Inference

**DOI:** 10.64898/2026.11.04.764238

**Authors:** Lingzhe Xu, Liu Zheng, Yangen Zhan, Yongbing Zhang, Kai Zhang

## Abstract

Spatially resolved protein abundance reveals cellular states and tissue organization that RNA expression cannot fully capture, yet direct spatial protein profiling remains costly and limited in coverage. Histology images routinely accompany spatial transcriptomics and offer complementary morphological information for protein inference.

Here, we introduce MorphoProt, a multimodal framework that jointly predicts spatial protein abundance from RNA expression, multiscale histology and tissue neighborhoods. MorphoProt uses RNA-guided cross-attention over five histological fields of view, so that the morphological context read for each spot is conditioned on its molecular state. The fused representation is refined by gated aggregation of RNA-derived information from physical and transcriptomic neighborhoods before joint protein prediction.

On held-out sections of human tonsil, mouse spleen and human liver, MorphoProt achieves mean protein-wise Spearman correlations of 0.652, 0.604 and 0.372, outperforming matched RNA-only and graph-based baselines and the histology-based HIPI and multimodal SPRINT baselines in all three tissues. Combining fields of view, rather than any single one, carries the histology gain: in spleen and liver, the multiscale model outperforms every single-field model, and its learned scale preference separates white-pulp from red-pulp compartments. Predicted proteins preserve protein-defined tissue domains and their marker identity, and recover pathologist-annotated germinal centers as accurately as measured proteins.

## 1 Introduction

Proteins mediate signaling, metabolism and interactions between cells and their surroundings, so measuring protein abundance in tissue provides a functional view of cellular states and tissue organization. Spatial transcriptomics (ST) profiles RNA while preserving tissue context [1], but transcript abundance reflects protein abundance only incompletely: differences in translation and protein turnover cause RNA–protein discor-dance [2]. Spatially resolved protein measurements are therefore an important complement to transcriptomic profiling [3, 4].

Direct spatial protein measurement remains experimentally demanding. Multiplexed imaging and paired RNA–protein assays extend molecular characterization, but require additional reagents, assay optimization and protein detection. SPOTS and spatial CITE-seq demonstrate the value of measuring selected protein panels alongside RNA [3, 4], yet their coverage is bounded by the assay and antibody panel. Consequently, many spatial transcriptomics datasets lack protein measurements, and computational prediction from paired reference data offers a way to estimate protein abundance in sections where proteins were not measured.

Methods developed for CITE-seq predict surface proteins from single-cell transcriptomes [5, 6, 7], but do not use spatial context. Existing spatial approaches infer proteins mainly from transcriptomic profiles and spatial relationships. DGAT learns spatially informed representations from paired RNA–protein data through modality-specific graphs and cross-modal supervision [8]. STProtein combines graph learning with multitask training [9], and DPAS-Graph models physical proximity and transcriptomic similarity as distinct relation channels for spatial RNA-to-protein prediction [10]. These studies establish transcriptomic information and spatial neighborhoods as informative inputs. Histology images add information on tissue morphology and architecture. Pathology foundation models [11] and paired resources such as HEST-1k [12] have made H&E an input for molecular prediction, from super-resolved and generated gene expression [13, 14] to joint transcriptomic–histology foundation models [15]. For proteins, HIPI predicts multiplexed protein levels from histology tiles [16], HEX predicts spatial protein expression from H&E images [17], TissueCraftAI generates spatial molecular maps from histology with a text-guided generative framework [18], and SPRINT combines histology with spatial transcriptomics for spatial protein reconstruction [19]. When RNA and proteins are both measured, SpatialGlue and DIME integrate them across modalities and serial sections [20, 21], and Steamboat uses multiscale attention to resolve cellular interactions [22]. Despite this progress, two questions lack joint modeling: how the molecular state of a spot can guide the selection of morphological information across scales, and how the distinct neighborhood relations represented by physical proximity and transcriptomic similarity can be separated and integrated.

Here, we present MorphoProt, a morphology-aware framework for spatial protein inference (Fig. 1). For each spot, the RNA profile guides cross-attention over five H&E fields of view, and two gated graph pathways add physical and transcriptomic neighborhood context before the protein panel is decoded jointly, so that the molecular state of a spot, its multiscale morphology and both neighborhood relations are modeled together. We evaluate MorphoProt on held-out sections of human tonsil, mouse spleen and human liver, against RNA-only, graph-based, histology-based and multimodal predictors. MorphoProt achieves the highest mean protein-wise Spearman correlation in all three tissues and improves on a matched RNA-only network by 0.038–0.065. Comparing the five-field model with single-field models shows that the contribution of multiscale morphology is tissue-dependent: in spleen and liver, the five-field model outperforms every single-field model.

**Figure 1:**
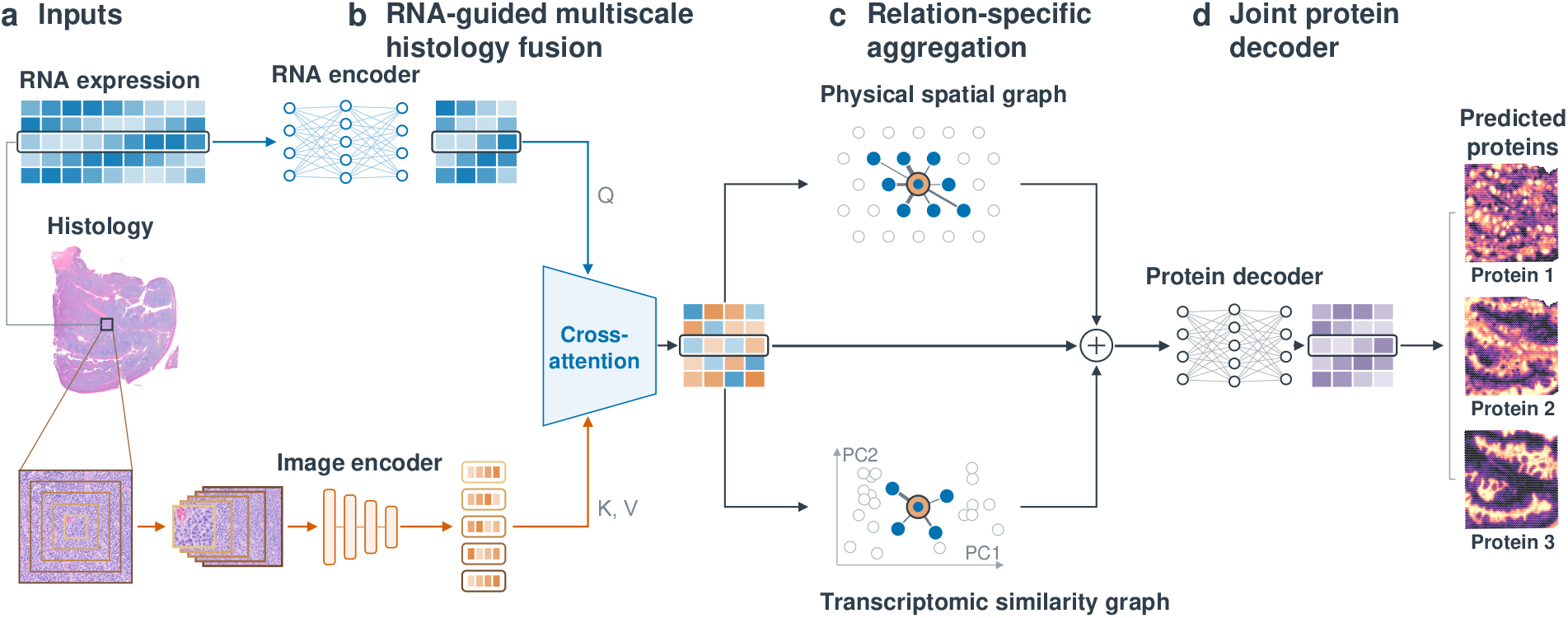
MorphoProt overview. **a**, Inputs per spot: the RNA profile and five H&E fields of view centered on the spot (tonsil 55–275 *µ*m; spleen and liver about 70–620 *µ*m). **b**, The RNA representation queries (Q) five tokens of a frozen image encoder (K, V; UNI2-H for tonsil, ResNet18 for spleen and liver) by cross-attention; the attended summary is added to it to form the fused feature. **c**, GATv2 blocks over a physical graph (eight nearest spots) and a transcriptomic graph (four nearest spots in RNA-PCA space): the fused center feature scores neighbors whose messages are their RNA representations, and gated messages are added to the fused feature. **d**, A joint decoder predicts the protein panel; maps show predicted proteins 1–3 (MS4A1, CD3E and KRT5) on the held-out tonsil section.

The learned scale preference follows the compartments of the spleen, with fine fields associated with white-pulp markers and coarse fields with red-pulp macrophage markers.

Beyond accuracy, domains derived from predicted tonsil proteins recover the five main regions defined by measured proteins with their marker profiles, and a germinal-center score from predicted proteins identifies pathologist-annotated germinal centers (AUPRC 0.786). MorphoProt thus improves spatial protein prediction while recovering tissue structures with clear biological meaning.

## 2 Results

### 2.1 Overview of MorphoProt

MorphoProt combines the RNA profile of each spot with five H&E fields of view and two neighborhood graphs (Fig. 1): the RNA representation weights the fields through cross-attention, the fused representation aggregates physical and transcriptomic neighbors through gated GATv2 blocks, and a joint decoder predicts the full protein panel (Section 3).

### 2.2 MorphoProt improves spatial protein prediction across three tissues

We evaluated Morphoprot on held-out sections of human tonsil, mouse spleen and human liver (4,908, 2,768 and 7,299 test spots; 29, 21 and 31 proteins; Section 3.1). For every method, predictions of models trained with seeds 42, 99, 7 and 1024 were averaged, and Spearman correlation (SCC) was computed for each protein across the test spots. The comparison includes a matched RNA-only network, the graph-based DGAT [8] and DPAS-Graph [10], the image-only HIPI [16] and the multimodal SPRINT [19], all trained on the same section partitions, protein panels and seed-specific training subsets (Section 3.4).

MorphoProt reached mean SCCs of 0.652, 0.604 and 0.372 and was the most accurate predictor in all three tissues (Table 1, Fig. 2a,b). It exceeded the RNA-only network by 0.038, 0.040 and 0.065, and SPRINT, the strongest multimodal baseline, by 0.018 (paired 95% CI over proteins [0.001, 0.037]), 0.068 ([0.052, 0.086]) and 0.018 ([™0.023, 0.057]), predicting 20/29, 21/21 and 21/31 proteins more accurately than SPRINT.

**Table 1:** Prediction performance on held-out sections. Mean protein-wise SCC and PCC, and pooled RMSE, of four-seed ensembles. Bold marks the best value.

|  | Tonsil |  |  | Spleen |  |  | Liver |  |  |
|---|---|---|---|---|---|---|---|---|---|
| Method | SCC ↑ | PCC ↑ | RMSE ↓ | SCC ↑ | PCC ↑ | RMSE ↓ | SCC ↑ | PCC ↑ | RMSE ↓ |
| RNA-only NN | 0.6136 | 0.6362 | **0.8077** | 0.5638 | 0.6382 | 1.2350 | 0.3064 | 0.3283 | 1.2300 |
| DGAT | 0.5996 | 0.6177 | 0.8748 | 0.5632 | 0.5833 | 1.2446 | 0.2930 | 0.2962 | 1.2696 |
| DPAS-Graph | 0.5778 | 0.5904 | 1.0594 | 0.5447 | 0.5336 | 1.2385 | 0.2884 | 0.2965 | 1.3312 |
| HIPI | 0.2497 | 0.2884 | 1.0409 | 0.2025 | 0.1938 | 1.4954 | 0.0160 | 0.0190 | 1.3770 |
| SPRINT | 0.6338 | 0.6500 | 0.8334 | 0.5362 | 0.6120 | 1.2496 | 0.3541 | 0.3373 | 1.2078 |
| MorphoProt | **0.6515** | **0.6740** | 0.8215 | **0.6041** | **0.6700** | **1.2345** | **0.3717** | **0.3855** | **1.1653** |

**Figure 2:**
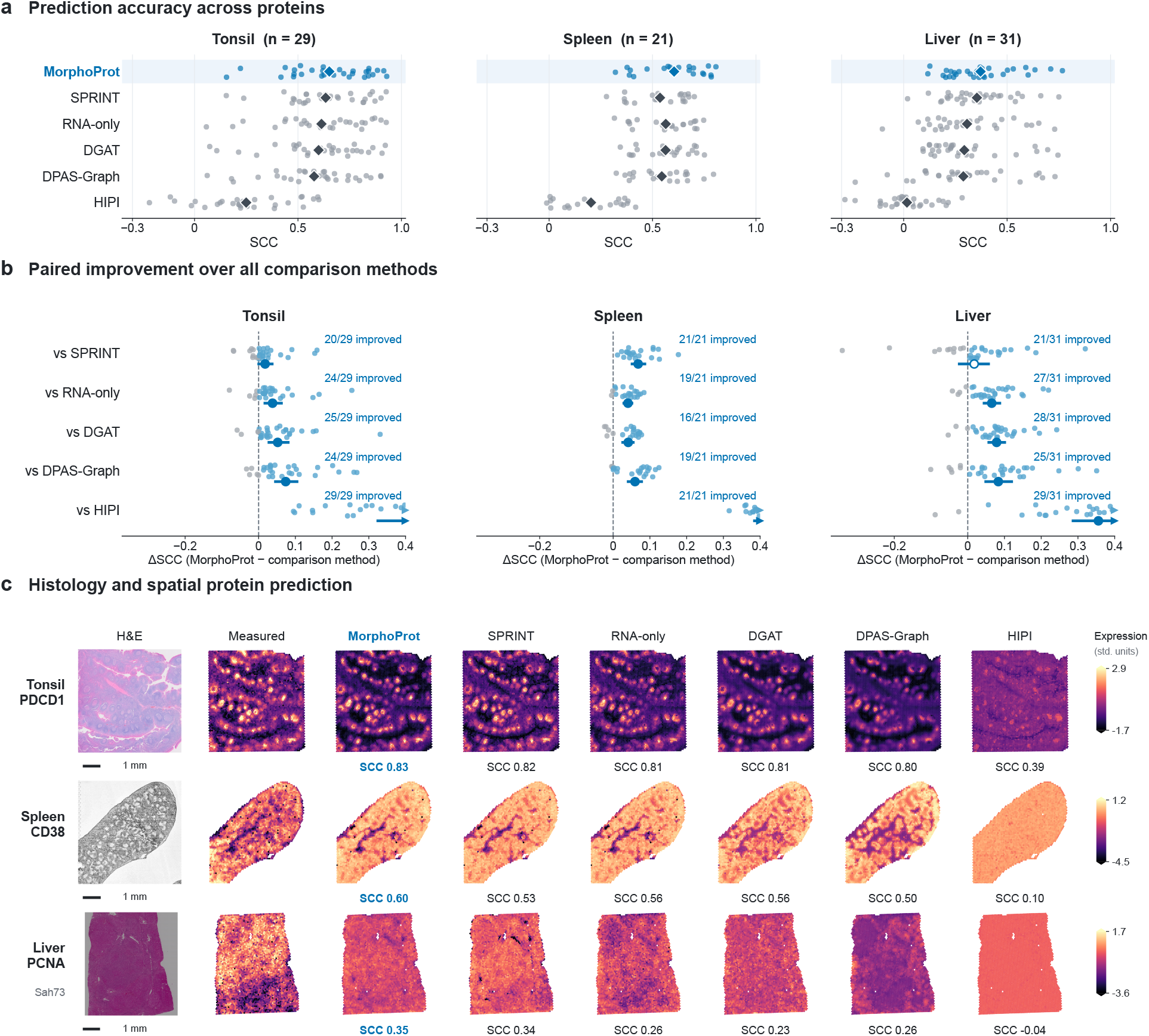
Prediction accuracy and spatial reconstruction across three tissues. **a**, Protein-wise SCC on held-out test spots for the six methods; dots are proteins and diamonds method means. **b**, Paired SCC difference between MorphoProt and each comparison method; circles and bars give the mean and 95% CI over proteins, counts give proteins with a positive difference, and arrowheads mark values beyond the axis. **c**, H&E, measured expression and the predictions of all six methods for one representative protein per tissue, with SCC over all test spots; maps in a row share one color range (1st–99th percentile of measured values), and liver maps show section Sah73. Scale bars, 1 mm.

The improvement is broad across proteins and largest for cell populations with a distinct morphology. Mor-phoProt improved over the RNA-only network for 24/29, 19/21 and 27/31 proteins (Fig. 2b; Supplementary Table S4), led by the plasma-cell marker SDC1 in tonsil (+0.254), the marginal-zone macrophage marker CD169 in spleen (+0.084) and the myeloid integrin ITGAM in liver (+0.220). For a representative protein per tissue (PDCD1, CD38 and PCNA), MorphoProt gave the highest SCC among the six predictors and reproduced the measured spatial patterns, such as the PDCD1-high germinal centers of the tonsil (Fig. 2c). Against the cognate transcripts of the same proteins, MorphoProt reached 0.660 and 0.604 on the matched tonsil and spleen panels, compared with 0.421 and 0.196, and was more accurate for 21/22 and 21/21 markers.

### 2.3 Combining fields of view helps where the fields carry complementary mor-phology

#### Modalities

Histology and neighborhoods contribute complementary information (Table 2). Adding image fusion to the RNA network raised SCC by 0.022, 0.022 and 0.051, adding the two neighborhood graphs by 0.029, 0.020 and 0.024, and combining both gave the best model in every tissue. In liver, histology contributed twice the gain of neighborhoods.

**Table 2:** Ablation study. Mean protein-wise SCC and PCC, and pooled RMSE, of four-seed ensembles. Bold marks the best value in each column. Single-scale uses the validation-selected field; protein-specific query denotes protein-specific attention (Section 3.4).

|  | Tonsil |  |  | Spleen |  |  | Liver |  |  |
|---|---|---|---|---|---|---|---|---|---|
| Ablation setting | SCC ↑ | PCC ↑ | RMSE ↓ | SCC ↑ | PCC ↑ | RMSE ↓ | SCC ↑ | PCC ↑ | RMSE ↓ |
| Full model | **0.652** | **0.674** | 0.821 | **0.604** | **0.670** | 1.234 | **0.372** | **0.386** | **1.165** |
| RNA-only | 0.614 | 0.636 | 0.808 | 0.564 | 0.638 | 1.235 | 0.306 | 0.328 | 1.230 |
| RNA + graphs | 0.642 | 0.659 | 0.810 | 0.584 | 0.653 | 1.217 | 0.331 | 0.345 | 1.189 |
| RNA + histology | 0.636 | 0.661 | 0.867 | 0.586 | 0.654 | 1.223 | 0.357 | 0.378 | 1.217 |
| Single-scale | 0.651 | 0.672 | **0.799** | 0.588 | 0.658 | 1.214 | 0.361 | 0.378 | 1.184 |
| No spatial graph | 0.642 | 0.666 | 0.865 | 0.590 | 0.660 | 1.230 | 0.360 | 0.380 | 1.197 |
| No transcriptomic graph | 0.649 | 0.672 | 0.823 | **0.604** | **0.670** | 1.193 | 0.367 | 0.381 | 1.177 |
| Token concat. | 0.650 | 0.673 | 0.879 | 0.594 | 0.655 | 1.300 | 0.369 | 0.382 | 1.174 |
| Protein-specific query | 0.649 | 0.673 | 0.838 | 0.598 | 0.663 | **1.185** | 0.371 | 0.381 | 1.171 |

#### Fields of view

To test whether the histology gain depends on combining scales, we retrained the full model with each single field of view (Fig. 3a). In spleen and liver, every single-field model fell below the five-field model, by 0.009–0.016 and 0.011–0.022. Relative to the validation-selected field, the five-field model gained 0.016 (95% CI [0.012, 0.020]) in spleen and 0.011 ([0.004, 0.017]) in liver, and improved 20/21 and 23/31 proteins (Supplementary Table S9). In tonsil, where histology adds 0.009 beyond RNA and neighborhoods, the validation-selected single field matched the five-field model (0.651 against 0.652).

**Figure 3:**
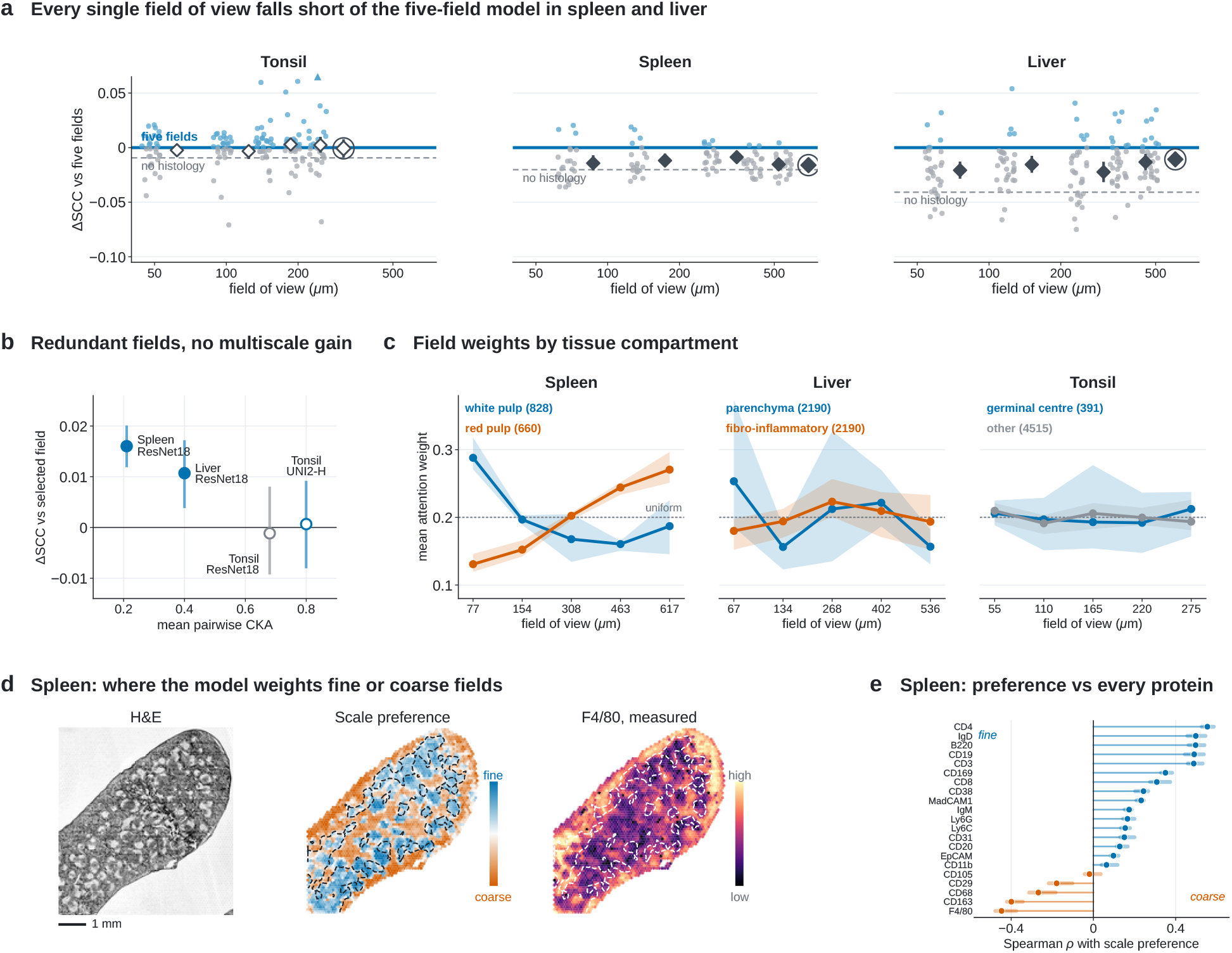
Multiscale histology with RNA-guided cross-attention. **a**, Per-protein SCC of the full model retrained with each single field of view minus that of the five-field model, against field size. Dots are proteins; diamonds and bars give the mean and 95% CI; the ringed diamond marks the validation-selected field and the dashed line the model without histology. **b**, Mean pairwise CKA between the five field embeddings of the test spots against the gain of the five-field model over the validation-selected field (mean and 95% CI over proteins; filled: CI excludes zero). **c**, Mean attention weight per field in each compartment (lines, mean of four seeds; bands, seed range; dotted line, uniform weight): spleen white and red pulp, liver parenchyma and fibro-inflammatory regions, tonsil germinal centers (pathologist annotation) and other spots; numbers of spots in parentheses. **d**, Spleen test section: H&E, per-spot scale preference (finest two minus coarsest two fields) and measured F4/80; outlines mark B220-high white pulp. Scale bar, 1 mm. **e**, Spearman correlation between scale preference and each measured spleen protein (mean and range over four seeds).

#### Field redundancy

The gain from combining fields tracked how much the fields differ (Fig. 3b). The five field embeddings of the test spots were least similar in spleen (mean pairwise CKA 0.21) and liver (0.40), where combining fields helped, and most similar in tonsil (0.80), where it did not. Replacing UNI2-H by ResNet18 in tonsil lowered this similarity to 0.68 but still left no multiscale gain (™0.001, [™0.009, 0.008]; Supplementary Tables S10, S11), so the absent gain in tonsil is not specific to the encoder.

#### Fusion and neighborhoods

RNA-guided cross-attention outperformed a token-concatenation Transformer over the same five fields, most clearly in spleen (+0.010, [0.005, 0.016]; 16/21 proteins), while using 1.32 million fewer parameters (Table 2; Supplementary Table S8). A single spot-level weighting of fields also outperformed protein-specific attention queries (spleen ™0.006, [™0.010, ™0.002]; Supplementary Table S12). Removing the physical graph lowered SCC by 0.010–0.014, more than removing the transcriptomic graph.

### 2.4 The RNA state of a spot determines which fields it reads

Because the RNA representation is the query, spots in different molecular states can weight the fields differently. In spleen, white-pulp spots placed most weight on the finest field (0.29, against 0.19 on the coarsest), whereas red-pulp spots placed most weight on the coarsest (0.27, against 0.13 on the finest), with the same ordering in all four seeds (Fig. 3c). Liver parenchyma also favored the finest field over fibro-inflammatory regions (0.25 against 0.18), with larger variation across seeds, and in tonsil, germinal-center and other spots both weighted all fields nearly uniformly (0.19–0.21), consistent with the redundant tonsil fields. The spleen pattern holds without compartment thresholds (Fig. 3d,e): spots that weight the finest fields are enriched for white-pulp markers (CD4, IgD, B220, CD19 and CD3; Spearman *ρ* = 0.49–0.55), whereas spots that weight the coarsest fields are enriched for red-pulp macrophage markers (F4/80, CD163 and CD68; *ρ* = ™0.45 to ™0.27), with the same sign in all four seeds (Supplementary Fig. S7, Table S13). Lymphocyte-dense white pulp is recognizable from local cell density, whereas the red-pulp macrophage network is defined by tissue architecture at larger scales.

### 2.5 Predicted proteins preserve tissue domains and germinal centers

Protein-defined tissue domains were recovered more faithfully from MorphoProt predictions than from any other predictor (Fig. 4a,c). Clustering measured and predicted proteins independently, MorphoProt reached an adjusted Rand index (ARI) of 0.703 in tonsil and 0.334 in spleen at *k* = 5, compared with 0.513 and 0.195 for SPRINT and 0.656 and 0.280 for RNA-only, and led both at 5/6 and 6/6 values of *k* = 3, …, 8. The predicted domains keep their biological identity (Fig. 4b): in tonsil, the five domains defined by measured proteins correspond to germinal centers, B-cell follicles, the T-cell zone, epithelium and stroma, and the domains defined by predicted proteins show the same marker profiles (correlation 0.95–0.98 per domain), with 86.8% of spots assigned to the same domain. The germinal-center domain of the predicted proteins contains 75% of the pathologist-annotated germinal-center spots (measured proteins, 73%).

**Figure 4:**
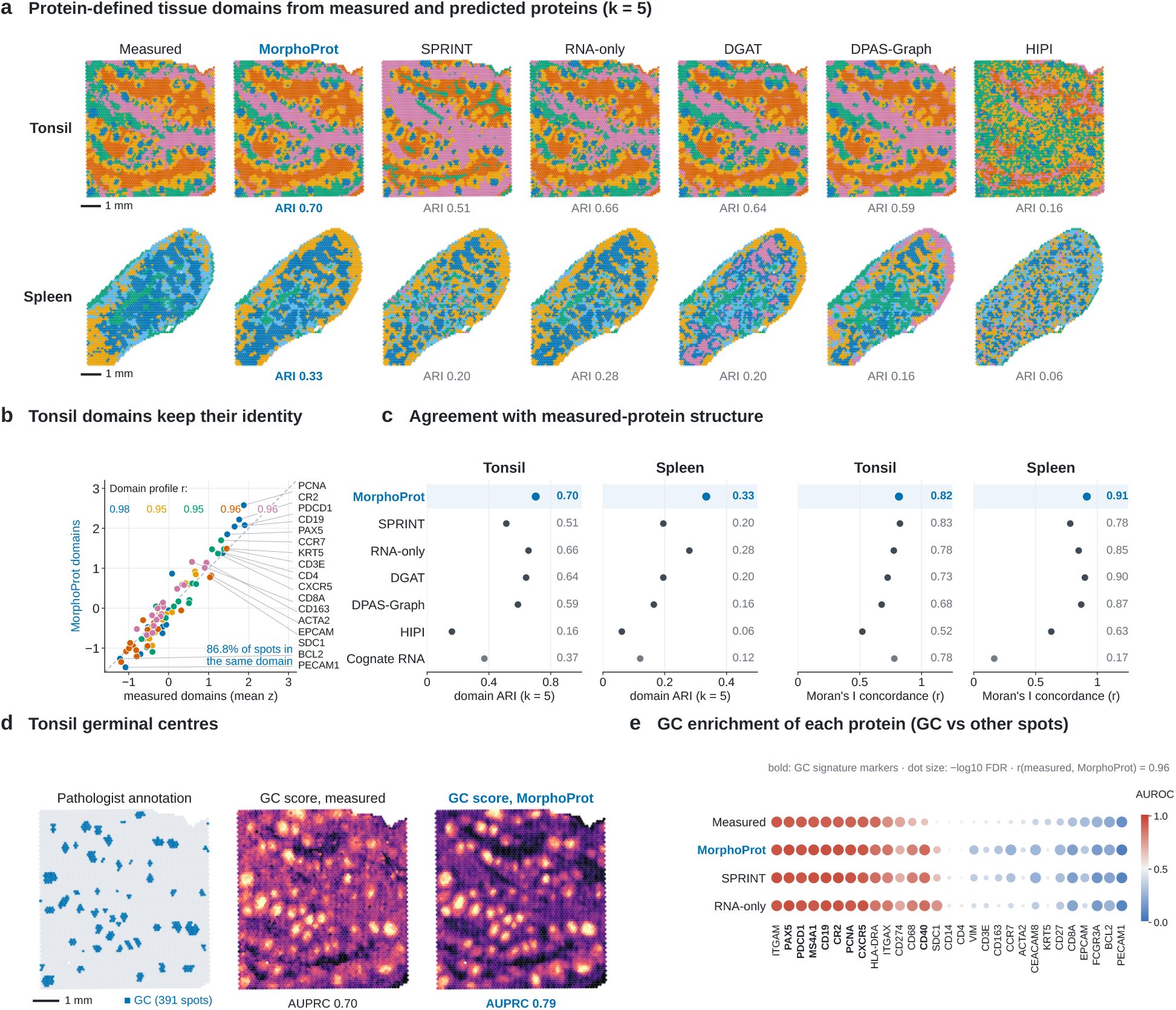
Predicted proteins preserve tissue organization. **a**, *k*-means domains (*k* = 5) of measured proteins and of each prediction on the matched tonsil and spleen panels, with ARI to the measured domains; predicted domains are colored by their best-matching measured domain. Scale bars, 1 mm. **b**, Mean standardized expression of 17 markers per tonsil domain, measured versus MorphoProt; *r*, per-domain correlation. **c**, Domain ARI (*k* = 5; bars, seed range) and correlation between predicted and measured bivariate Moran’s I matrices. **d**, Pathologist germinal-center (GC) annotation of the held-out tonsil section and the eight-marker GC score from measured and MorphoProt proteins, with the AUPRC of the score. **e**, Enrichment of each protein in GCs (AUROC of GC versus other spots; dot size, ™log_10_ BH-adjusted FDR) for measured proteins and three predictors; bold, GC-score markers.

Predicted proteins also preserve pairwise spatial co-localization: their bivariate Moran’s I matrix correlated with the measured matrix at 0.82 in tonsil and 0.91 in spleen (Fig. 4c).

Finally, predicted proteins identify pathologist-annotated germinal centers (Fig. 4d). A score averaging eight germinal-center markers reached an area under the precision–recall curve (AUPRC) of 0.786 with Mor-phoProt predictions, 0.702 with measured proteins and 0.398 with the corresponding mRNA; MorphoProt and SPRINT performed comparably (Supplementary Table S15). Protein by protein, enrichment in germinal centers from MorphoProt predictions agreed with that from measured proteins (correlation of AUROCs 0.96 across the 29 proteins; Fig. 4e), with the canonical germinal-center markers most enriched in both.

## 3 Methods

For *N* spots, let *X* ∈ ℝ^*N ×G*^ denote RNA features, *C* ∈ ℝ^*N ×*2^ spatial coordinates, and *Y* ∈ ℝ^*N ×P*^ the measured protein panel. The set *I* = {*I*^(1)^, …, *I*^(*L*)^} contains spot-centered images at *L* = 5 fields of view. Frozen encoders map these images to features 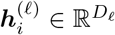. We learn *Ŷ* = *f*_*θ*_(*X, C, I*) from paired training sections and evaluate on held-out sections. At test time, RNA, images, coordinates, and test–test edges are available; protein labels are not inputs to the predictor. This defines a transductive prediction setting.

### 3.1 Datasets and Preprocessing

Tonsil contains a female section (4,194 spots) and a male section (4,908), with 29 targets after excluding four control antibodies; the primary direction is female to male. Mouse spleen uses SPOTS GSE198353 [3], with 2,568 training and 2,768 test spots and 21 proteins. Human liver GSE278662 [23] uses N123/N126 for training (6,714 spots) and Sah73/Sah80 for testing (7,299), with 31 proteins.

The primary RNA pipeline filters genes detected in fewer than three training spots and selects 3,500 genes using training-section variability with retained protein-marker genes. Normalization to 10,000 counts and log transformation support feature selection; model inputs use log-transformed raw RNA and training-section scaling. Spleen/liver proteins use training-fitted scaling of log(1 + *x*). In contrast, tonsil protein matrices were standardized separately within each section. Liver missing image features are zero-filled (1, 3, 8, 14, and 26 patches across ascending scales).

### 3.2 Multiscale Histology Encoding and RNA-Guided Fusion

#### Encoding RNA

For spot *i*, the RNA encoder maps the 3,500-gene vector ***x***_*i*_ to 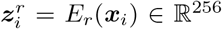 through two fully connected layers (widths 768 and 256), each followed by batch normalization, GELU and dropout, and three residual MLP blocks that add their update to the input with weight 0.5. The same encoder supplies the representations of center spots and their neighbors.

#### Encoding multiscale histology

Five spot-centered fields of view are encoded by a frozen image encoder, with patches resized to the 224-pixel encoder input. In tonsil, 256–1,280-pixel fields at 0.215 *µ*m per pixel (55–275 *µ*m, one to five spot diameters) are encoded by the pathology foundation model UNI2-H [11, 24] (1,536 dimensions). In spleen and liver, whose images have about 4–5 *µ*m per pixel, 16–128-pixel fields (about 70–620 *µ*m) are encoded by ImageNet ResNet18 [25] (512 dimensions). A scale-specific projection maps each feature 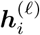 to a token

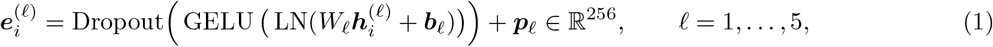

where LN is layer normalization and ***p***_*l*_ a learned scale embedding.

#### RNA-conditioned integration of image context

The RNA representation is a single query over the five image tokens. For heads *h* = 1, …, 4 of width *d*_*h*_ = 64, affine maps *Q*_*h*_, *K*_*h*_ and *V*_*h*_ form

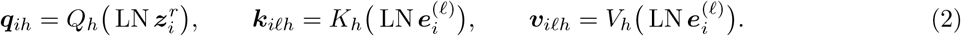

The weight of field *l* and the attended image summary are

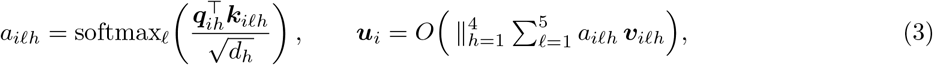

where *O* projects the concatenated heads back to 256 dimensions. The weights over fields thus depend on the transcriptional state of the spot. The image summary enters through a learned scalar and a residual post-fusion map,

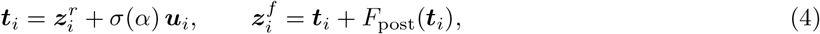

with *σ*(*α*) initialized to 0.10 and *F*_post_ an LN–linear–GELU–dropout map; RNA thus remains the backbone of the representation and histology refines it.

### 3.3 Neighborhood Aggregation and Protein Decoding

#### Dual neighborhoods

The physical neighborhood *N*_*s*_(*i*) contains the eight nearest spots in tissue coordinates and the transcriptomic neighborhood *N*_*f*_ (*i*) the four nearest spots in a 64-dimensional RNA-PCA space fitted on the inner training partition and applied to validation and test RNA. Neighbors are searched within each partition, excluding the spot itself. For relation *g* ∈ {*s, f*}, the edge descriptor 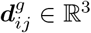 holds the first two neighbor-minus-center components and the Euclidean distance in the corresponding coordinate or PCA space.

#### Gated GATv2 aggregation

The center contributes its fused representation 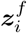, whereas each neighbor contributes 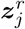: histology shapes how incoming messages are weighted, while the messages remain RNA-derived. Each relation uses a four-head GATv2 block [26] with score and attention weight

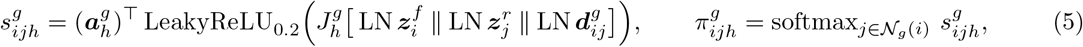

and the aggregated message is

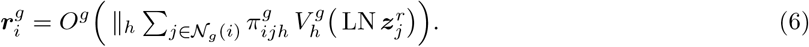

Each relation enters as a scaled, feature-wise gated residual update,

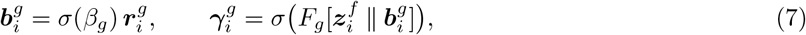

with *σ*(*β*_*g*_) initialized to 0.08 and *F*_*g*_ a linear–GELU–dropout–linear map, and the decoder input is

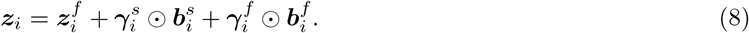

#### Joint protein decoding

A decoder (256-unit linear layer with batch normalization, GELU and dropout, a residual MLP block and a linear output) maps ***z***_*i*_ to ***o***_*i*_ ∈ ℝ^*P*^, and the prediction ***ŷ***_*i*_ = ***o***_*i*_ + 0.15 *A****o***_*i*_ couples proteins, where the trainable *A* ∈ ℝ^*P ×P*^ is initialized from Pearson correlations among training-target proteins.

#### Loss

Training combines squared error, concordance and pairwise rank agreement. For a minibatch of *B* spots, learned protein-specific log variances *s*_*p*_ (initialized to zero, clamped to [™5, 5]) weight the squared errors,

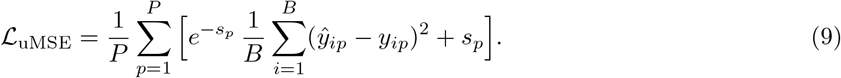

The concordance term uses Lin’s concordance correlation [27] averaged over proteins, and the ranking term a set Q of 1,024 untied spot pairs sampled per batch and shared across proteins:

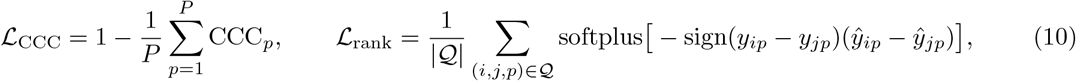

and the objective is L = L_uMSE_ + 0.15 L_CCC_ + 0.05 L_rank_.

### 3.4 Training and Evaluation

#### Optimization

MorphoProt and all neural controls are trained with seeds 42, 99, 7 and 1024, each reserving 15% of training-section spots for validation. AdamW (learning rate 4 × 10^*™*4^, weight decay 10^*™*4^, batch size 128) is used with gradient clipping at norm 5 and a plateau scheduler that halves the learning rate after four epochs without improvement. Training runs for at most 40 epochs in tonsil and spleen (early-stopping patience eight) and 160 epochs in liver (patience 30), and the checkpoint is chosen by validation mean SCC. Dropout is 0.15 in the RNA encoder, 0.12 in the image, attention, graph and decoder blocks, and 0.075 in the gates. Liver image features are standardized per dimension with statistics of the inner-training subset of each seed, and liver coordinates are offset to separate sections. Ensemble predictions are the arithmetic mean of the four selected models.

#### Controls and comparison methods

Four neural variants separate the modalities: RNA-only feeds the RNA encoder directly to the decoder; RNA+graphs adds both neighborhood pathways to that representation; RNA+histology keeps the full fusion backbone and omits graph aggregation; the full model uses both. Branch-removal controls delete the physical or transcriptomic graph block, its gate and its residual coefficient. The fusion control passes one RNA token and the five image tokens through two Transformer encoder layers and keeps the graph modules and decoder. Single-field models use one field of view as the only image token (five fields per tissue); the validation-selected field has the highest mean validation SCC over seeds. Protein-specific queries add one learned query per protein (best of three variants). Every control is retrained with the same seeds, inputs and settings as the full model. DGAT [8], HIPI [16] and SPRINT [19] are trained from their official code, and DPAS-Graph [10] from code released by an independent validation study with its graph builder re-implemented from the paper. All use the same section partitions, protein panels and seed-specific 85% training subsets; validation and checkpoint selection use only the withheld training-section spots, and no test labels enter fitting or selection (Supplementary Methods).

#### Metrics and downstream analyses

The primary metric is the mean protein-wise Spearman correlation over the test spots, 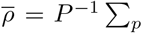 corr(rank(*Y*_:*p*_), rank(*Ŷ*_:*p*_)), with average ranks for ties; PCC and pooled RMSE supplement it. Confidence intervals for paired model contrasts use 100,000 paired protein resamples; in the figures, filled symbols mark intervals that exclude zero. Scale preference is the attention weight on the two finest minus the two coarsest fields, averaged over heads, computed on test spots for each of the four checkpoints. Compartment field weights average the head-mean attention over the test spots of each compartment: spleen white pulp is the top 30% of measured B220 and red pulp the top 30% of F4/80 among the remaining spots, each averaged over the spot and its six nearest neighbors; liver fibro-inflammatory and parenchymal spots are the top and bottom 30% of the neighbor-averaged mean of standardized ACTA2, VIM and PTPRC; tonsil spots are split by the pathologist annotation. Field redundancy is the mean pairwise CKA between the five field embeddings of the test spots. For domain analyses, measured proteins, each prediction and cognate-RNA proxies on matched panels are standardized and clustered independently by *k*-means (*k* = 3, …, 8, seeds 0–9, 20 initializations); agreement with measured-protein clusters is the ARI averaged over seeds; and predicted domains are matched to measured domains by Hungarian assignment. Tonsil domains are named by the strongest of five marker groups (Supplementary Methods). Bivariate Moran’s I uses row-standardized six-nearest-neighbor weights, and concordance is the Pearson correlation of its off-diagonal entries with the measured-protein matrix. Germinal centers (GC) of the held-out tonsil section are taken from the pathologist annotation released with DGAT [8]; in each clustering (*k*-means, and Leiden as a check), the cluster with the highest F1 against it is called GC (Supplementary Table S16). The GC score is the mean standardized value of eight canonical GC markers (PAX5, CD19, MS4A1, CXCR5, CR2, CD40, PDCD1 and PCNA), scored by AUPRC (prevalence 0.080) with 95% CIs from a spatial block bootstrap (74 blocks of about 0.8 mm, 2,000 resamples). Per-protein GC enrichment is the AUROC of GC versus other spots, with Benjamini–Hochberg-adjusted Wilcoxon rank-sum FDR.

## 4 Discussion

MorphoProt integrates RNA-conditioned multiscale histology with RNA-derived neighborhood information for spatial protein prediction. Across three tissues it achieved the highest mean protein-wise SCC among the evaluated methods and preserved protein-defined tissue domains, marker profiles, spatial co-localization and germinal-center structures. Combining fields of view improved prediction where the fields were complementary, and the learned field weights differed between tissue compartments, most clearly between white and red pulp in spleen.

The contributions of histology and multiscale integration varied across tissues. The tonsil fields were largely redundant under both encoders, and the compartment-specific weighting of fields was strongest in spleen, where the fields were least redundant. These findings indicate that the benefit of multiscale integration depends on tissue context and that a universal physical field size may not be optimal across datasets.

Several limitations remain. Each tissue was evaluated using a single held-out partition, and the current confidence intervals do not capture variability across independent donors or tissue sections. This reflects the few available datasets with paired spatial transcriptomic and proteomic measurements. Protein-wise comparisons were not adjusted for multiple testing. Although MorphoProt achieved higher mean SCC than SPRINT in all three tissues, the margin was small in tonsil and the paired interval included zero in liver, and the two methods showed comparable performance in germinal-center identification. Generalization across donors, staining protocols, platforms and unseen protein panels remains to be established; future work should prioritize independent validation and investigate pathology foundation-model features and adaptive field selection based on tissue geometry and image resolution. The RNA-conditioned integration of multiscale context may also extend to other spot-level prediction tasks in spatial omics.

## Data and Code Availability

Code is available at https://github.com/xlzhhh123/MorphoProt. The liver and spleen data are available from GEO (GSE278662, GSE198353); the tonsil data are the 10x Genomics CytAssist FFPE Human Tonsil (Protein Expression) datasets.

## References

[1] Ståhl, P. L. et al. Visualization and analysis of gene expression in tissue sections by spatial transcriptomics. Science 353, 78–82 (2016).

[2] Vogel, C. & Marcotte, E. M. Insights into the regulation of protein abundance from proteomic and transcriptomic analyses. Nature Reviews Genetics 13, 227–232 (2012).

[3] Ben-Chetrit, N. et al. Integrated protein and transcriptome spatial profiling. Nature Biotechnology 41, 788–793 (2023).

[4] Liu, Y. et al. High-plex protein and whole transcriptome co-mapping at cellular resolution with spatial CITE-seq. Nature Biotechnology 41, 1405–1409 (2023).

[5] Zhou, Z., Ye, C., Wang, J. & Zhang, N. R. Surface protein imputation from single cell transcriptomes by deep neural networks. Nature Communications 11, 651 (2020).

[6] Lakkis, J. et al. A multi-use deep learning method for CITE-seq and single-cell RNA-seq data integration with cell surface protein prediction and imputation. Nature Machine Intelligence 4, 940–952 (2022).

[7] Hanhart, D., Gossi, F., Rapsomaniki, M. A., Kruithof-de Julio, M. & Chouvardas, P. ScLinear predicts protein abundance at single-cell resolution. Communications Biology 7, 267 (2024).

[8] Wang, H. et al. DGAT: a dual-graph attention network for inferring spatial protein landscapes from transcriptomics. Nature Communications 17, 6702 (2026).

[9] Jiang, Z., Yuan, Y., Hu, L. & Pang, W. STProtein: predicting spatial protein expression from multi-omics data. arXiv preprint arXiv:2602.05811 (2026).

[10] Xu, M., Dong, Z. & Xia, B. DPAS-Graph: adaptive spatial-feature relation learning for spatial RNA-to-protein prediction and virtual protein profiling. Briefings in Bioinformatics 27, bbag480 (2026).

[11] Chen, R. J. et al. Towards a general-purpose foundation model for computational pathology. Nature Medicine 30, 850–862 (2024).

[12] Jaume, G. et al. HEST-1k: A dataset for spatial transcriptomics and histology image analysis. In Advances in Neural Information Processing Systems (Datasets and Benchmarks Track) (2024).

[13] Zhang, D., Schroeder, A., Yan, H. et al. Inferring super-resolution tissue architecture by integrating spatial transcriptomics with histology. Nature Biotechnology 42, 1372–1377 (2024).

[14] Wood, R., Hu, Y., Rittscher, J. & Li, B. GenST: A generative cross-modal model for predicting spatial transcriptomics from histology images. In Proceedings of the MICCAI Workshop on Computational Pathology, vol. 316 of Proceedings of Machine Learning Research, 52–65 (2026).

[15] Xiang, J., Hou, S., Li, Y. et al. A multimodal foundation model of spatial transcriptomics and histology for biological discovery and clinical prediction. arXiv preprint arXiv:2604.03630 (2026).

[16] Zeira, R., Anavy, L., Yakhini, Z., Rivlin, E. & Freedman, D. HIPI: Spatially resolved multiplexed protein expression inferred from H&E WSIs. PLOS Computational Biology 20, e1012501 (2024).

[17] Li, Z., Li, Y., Xiang, J. et al. AI-enabled virtual spatial proteomics from histopathology for interpretable biomarker discovery in lung cancer. Nature Medicine 32, 231–244 (2026).

[18] Pang, M., Roy, T. K., Wu, X. & Tan, K. Query-driven generative AI synthesizes multi-modal spatial omics from histology. bioRxiv (2025).

[19] Wang, Y., Sun, Y., Liu, X. & Zhao, T. SPRINT enables spatial proteomic profile reconstruction through multimodal fusion of histology and transcriptomics. Research Square preprint (2026).

[20] Long, Y., Ang, K. S., Sethi, R. et al. Deciphering spatial domains from spatial multi-omics with SpatialGlue. Nature Methods 21, 1658–1667 (2024).

[21] Sun, P., Mou, T., Huang, X. & Zheng, X. Achieving spatial multiomics diagonal integration from unaligned serial sections with DIME. Genome Research 36 (2026).

[22] Liang, S., Tang, J., Wang, G. & Ma, J. Steamboat: Attention-based multiscale delineation of cellular interactions in tissues. bioRxiv (2025).

[23] Gripshover, T. C. et al. Visium spatial transcriptomics and proteomics identifies novel hepatic cell populations and transcriptomic signatures of alcohol-associated hepatitis. Alcohol: Clinical and Experimental Research 49, 106–116 (2025).

[24] Mahmood Lab. UNI2-h: pretrained histopathology image encoder. https://huggingface.co/ MahmoodLab/UNI2-h. Model card; accessed 21 October 2026.

[25] He, K., Zhang, X., Ren, S. & Sun, J. Deep residual learning for image recognition. In Proceedings of the IEEE Conference on Computer Vision and Pattern Recognition, 770–778 (2016).

[26] Brody, S., Alon, U. & Yahav, E. How attentive are graph attention networks? In International Conference on Learning Representations (2022).

[27] Lin, L. I.-K. A concordance correlation coefficient to evaluate reproducibility. Biometrics 45, 255–268 (1989).

