## Supplementary Information for "MorphoProt: RNA-Conditioned Integration of Multiscale Histology for Spatial Protein Inference"

### MorphoProt: Supplementary Material

Every primary and ablation ensemble averages predictions from seeds 42, 99, 7 and 1024 before computing protein-wise correlations. RMSE is in tissue-specific standardized target units.

**Contents.** Supplementary Methods. Tables S1–S8: prediction performance, baseline seeds, matched controls, complete protein panels, cognate-RNA panels, liver sections, regression controls and fusion parameters. Tables S9–S16: single-field models, image-encoder control, field-of-view sensitivity, protein-specific queries, spleen scale preference for all proteins, downstream analyses of the multiscale controls, germinal-center AUPRC with spatial-block CIs and cluster-based germinal-center recovery. Figures S1–S8: liver controls, baseline seeds, complete-panel contrasts, cognate RNA, clustering NMI, additional spatial cases, attention over fields and the image-encoder control.

### Supplementary Methods

These details complete Sections 3.2–3.4 of the main text.

**Architecture details.** Each of the two fully connected RNA-encoder layers is followed by batch normalization, GELU and dropout; each of the three residual MLP blocks applies batch normalization and GELU before each of two linear maps and adds its update to the input with weight 0.5. Image patches are resized to the 224-pixel encoder input. Each scale-specific image projection is $\mathrm{Dropout}(\mathrm{GELU}(\mathrm{LN}(W_\ell h^{(\ell)} + b_\ell))) + p_\ell$, with $p_\ell$ a learned scale embedding. Cross-attention and graph blocks use attention and output dropout during training. The post-fusion map $F_{\text{post}}$ is LN–linear–GELU–dropout. In each GATv2 block, center, neighbor and edge features are layer-normalized separately, concatenated and projected before scoring; the LeakyReLU slope is 0.2. Neighbor searches discard the first returned neighbor (the spot itself), and the RNA PCA fitted on the inner training partition is applied to validation and test RNA. The gate networks $F_g$ are linear–GELU–dropout–linear. The decoder is a 256-unit linear layer with batch normalization, GELU and dropout, a residual MLP block, and a linear output layer. Protein log variances $s_p$ are initialized to zero and clamped to $[-5, 5]$. CCC uses population batch variances and covariance with denominator stabilization $10^{-6}$. The 1,024 ranking pairs are sampled with replacement and shared across targets; tied target pairs are excluded, and the ranking term is zero when none remain.

**Optimization.** Gradients are clipped at norm 5, and a plateau scheduler halves the learning rate after four epochs without improvement. Early-stopping patience is eight epochs in tonsil and spleen and 30 in liver. The RNA encoder and decoder input use dropout 0.15; image projections, cross-attention, post-fusion, graph blocks and the decoder residual block use 0.12; graph gates use 0.075. Liver coordinates are offset to separate sections; the transcriptomic graph can connect sections within a partition.

**Controls.** RNA-only feeds the RNA encoder directly to the decoder; RNA+graph adds both neighborhood pathways to that representation. Both omit image attention and the post-fusion map, whereas RNA+image keeps the full fusion backbone and omits graph aggregation. Ridge-regression controls on RNA, RNA principal components, and principal components with spatial-neighbor means use validation-selected regularization (details below). In the concatenation control, one RNA token and five image tokens are concatenated along the token axis with modality and position embeddings and passed through two Transformer encoder layers (four heads, feed-forward expansion four); the RNA-token output is normalized and passed through a residual post-fusion map, and the graph modules and decoder are retained (Table S8). Branch-removal controls delete the physical or transcriptomic graph block, its gate and its residual coefficient, and retrain with the same seeds and settings. Protein-specific queries add one learned query per protein, as a residual to the shared query, as a zero-initialized residual, or as its replacement; the best variant is reported.

**Official-code comparisons.** We evaluate DGAT on all three tissues using source commit `b0128f342de7`. The adapters use the original fixed RNA arrays, coordinates, protein panels, and 85% seed-specific labeled training subsets; the remaining 15% is withheld. DGAT retains its original training routine, sample-local RNA graph preprocessing, maximum 100 epochs, and evidence-based stopping. Its test graph uses RNA and coordinates; the RNA encoder feeds the trained protein decoder without test protein inputs. Protein outputs are restored to the common panel order.

DPAS-Graph has no author repository; we used the encoder and decoders released with an independent external-validation study (`zksdu/dpas-graph-external-validation`, commit `a6ab4dae511c`) with its published training settings (100 epochs, 8×SmoothL1 protein loss plus RNA reconstruction). Its graph builder is not public and was re-implemented from the paper: union of spatial ($k = 6$) and RNA-PCA ($k = 10$, 50 components fitted on the labeled training subset) neighbor edges with Gaussian two-channel weights, built per section. The upstream code selects epochs by test RMSE; we apply the same criterion to the withheld 15% validation spots. Every method is evaluated as the average of predictions from the same four seeds, and no test labels enter model fitting or checkpoint selection. DGAT and DPAS-Graph run with PyTorch 2.11.0+cu128 on an RTX 5070 Ti.

**Histology-based comparisons.** HIPI uses its official frozen pathology DINO-p8 encoder and regression MLP; SPRINT uses its official A2 architecture for tonsil and compact C2 for the low-resolution spleen/liver inputs. Both are trained on the common protein panels using the same seed-specific 85% labeled subsets. Their optimization and image handling are specified under *Reproduced baselines and alignment* below. Validation and checkpoint selection use only the withheld training-section spots, rather than the external test section used as validation in the SPRINT example notebooks. HIPI and SPRINT run with PyTorch 2.8.0+cu128 on the same RTX 5070 Ti. No pretrained protein-prediction checkpoint is used.

**Evaluation details.** Spearman correlation uses average ranks for ties; all evaluated targets have nonconstant values. Protein resampling uses analysis seed 20261017. RNA-proxy comparisons use only shared evaluable targets and the primary ensemble. $k$-means uses 20 initializations, and each clustering is also compared with the measured-protein clustering at the same $k$ and seed by NMI, AMI, FMI and homogeneity. A smoothed proxy averages each spot's cognate RNA with its six nearest spatial neighbors. Bivariate Moran's I is computed on standardized features and symmetrized. Domain maps show seed 0 at $k = 5$, and Hungarian relabeling is for display only. The Spearman correlation between scale preference and each measured protein is computed per seed and summarized by the mean and range. White pulp in main Fig. 3d is outlined as the top 30% of measured B220 averaged over each spot and its six nearest neighbors.

**Germinal-center evaluation.** The pathologist GC annotation of the held-out tonsil section was taken from the processed DGAT data release (`Tonsil_AddOns_ADT.h5ad`, `obs['pathology']`) (Wang et al.) and matched to our spots by barcode (all 4,908 spots; two without a label excluded). Each representation (measured proteins, four-seed ensemble predictions on the 29 targets, the cognate-RNA proxy on 22 genes, and the 3,500 input RNA features reduced to 30 principal components) was standardized and clustered by $k$-means ($k = 3, \ldots, 8$, seeds 0–9, 20 initializations) and, as a check, by Leiden clustering with the resolution tuned to $k = 5$. For each clustering, the cluster with the highest F1 against the annotation was called GC, and F1, precision, recall, MCC, Jaccard index, Cohen's $\kappa$, balanced accuracy, accuracy and specificity were computed; this selection step is applied identically to every representation. The cluster-free GC signature is the mean standardized value of PAX5, CD19, MS4A1, CXCR5, CR2, CD40, PDCD1 and PCNA, the canonical GC markers named by DGAT, scored by AUROC and AUPRC; MS4A1 is absent from the input RNA features and the proxy panel. Per-protein enrichment uses two-sided Wilcoxon rank-sum tests of GC versus other spots with Benjamini–Hochberg correction. Confidence intervals for AUPRC use a spatial block bootstrap: the section is divided into 74 blocks of about 0.8 mm (eight spot pitches), and blocks are resampled with replacement 2,000 times (seed 20261017). GC prevalence, the AUPRC of a random score, is 0.080.

**Domain identity.** For the tonsil domains at $k = 5$ (seed 0), the mean standardized expression of 17 markers in five groups (germinal center: CR2, PCNA, PDCD1; B follicle: CD19, PAX5, CXCR5, BCL2; T zone: CD3E, CD4, CD8A, CCR7; epithelium: KRT5, EPCAM, SDC1; stroma and vessels: ACTA2, PECAM1, CD163) is computed for measured domains from measured proteins and for MorphoProt domains, matched to measured domains by Hungarian assignment, from predicted proteins. Each measured domain is named by the group with the highest mean; the five domains receive five distinct names.

**Further evaluation details.**

**Matched liver protocol.** The full liver configuration uses per-dimension z-scoring of frozen ResNet18 image embeddings, fitted on each seed's inner-training subset. All six liver controls (four seeds each) use the same split, optimizer settings, maximum 160 epochs and patience 30. Checkpoints use validation mean SCC. Validation neighborhoods may use RNA/coordinates from all training sections; validation protein labels are withheld from fitting.

**Regression and proxy specification.** Ridge controls use validation-selected regularization. RNA ridge adds an inner-training scaler; PCA ridge uses 64 inner-training components. PCA+spatial ridge adds eight-neighbor means, with neighborhoods built within each partition. Penalties span $10^{-2}$ to $10^{6}$. RNA proxies are descriptive cognate-gene signals; their RMSE is omitted because their units are not calibrated to proteins. B220/CD45R uses total Ptprc. Liver has gene-level concordance statistics but no spot-level proxy panel in the downstream figures.

**Reproduced baselines and alignment.** HIPI and SPRINT use the official implementations adapted to the common spot-level panels. HIPI freezes the published pathology DINO-p8 encoder and trains its original MLP with MSE for 200 epochs, using Adam at $5.12 \times 10^{-4}$, batch 512 and validation-MSE checkpoint selection. Eight fixed official spatial/stain augmentation views per training spot are cached; validation and test are unaugmented. SPRINT uses A2 for tonsil and compact C2 for spleen/liver, Adam at $10^{-4}$ for 30 epochs, the original half-MSE/half-PCC loss, and validation-PCC checkpoint selection. All baseline validation subsets come from the same labeled training sections; no pretrained protein-prediction checkpoint is used. All methods retain common test proteins and spot ordering.

**Reproduction package.** Code is available at `https://github.com/xlzhhh123/MorphoProt`.

#### Supplementary Tables

**Table S1:** Primary four-seed prediction ensembles and liver section results. SCC/PCC are protein-wise means; RMSE is pooled over spot–protein entries. Section scores use the same predictions without refitting.

| Tissue / section | Method | SCC | PCC | RMSE |
|---|---|---|---|---|
| tonsil | MorphoProt | 0.6515 | 0.6740 | 0.8215 |
| tonsil | RNA-only NN | 0.6136 | 0.6362 | 0.8077 |
| tonsil | DGAT | 0.5996 | 0.6177 | 0.8748 |
| tonsil | DPAS-Graph | 0.5778 | 0.5904 | 1.0594 |
| tonsil | HIPI | 0.2497 | 0.2884 | 1.0409 |
| tonsil | SPRINT | 0.6338 | 0.6500 | 0.8334 |
| spleen | MorphoProt | 0.6041 | 0.6700 | 1.2345 |
| spleen | RNA-only NN | 0.5638 | 0.6382 | 1.2350 |
| spleen | DGAT | 0.5632 | 0.5833 | 1.2446 |
| spleen | DPAS-Graph | 0.5447 | 0.5336 | 1.2385 |
| spleen | HIPI | 0.2025 | 0.1938 | 1.4954 |
| spleen | SPRINT | 0.5362 | 0.6120 | 1.2496 |
| liver | MorphoProt | 0.3717 | 0.3855 | 1.1653 |
| liver | RNA-only NN | 0.3064 | 0.3283 | 1.2300 |
| liver | DGAT | 0.2930 | 0.2962 | 1.2696 |
| liver | DPAS-Graph | 0.2884 | 0.2965 | 1.3312 |
| liver | HIPI | 0.0160 | 0.0190 | 1.3770 |
| liver | SPRINT | 0.3541 | 0.3373 | 1.2078 |
| liver / Sah73 | MorphoProt | 0.3918 | 0.3898 | 1.3086 |
| liver / Sah80 | MorphoProt | 0.3660 | 0.3853 | 0.9698 |
| liver / Sah73 | RNA-only NN | 0.3236 | 0.3292 | 1.3732 |
| liver / Sah80 | RNA-only NN | 0.3322 | 0.3316 | 1.0363 |
| liver / Sah73 | DGAT | 0.3165 | 0.3106 | 1.4001 |
| liver / Sah80 | DGAT | 0.2552 | 0.2562 | 1.0961 |
| liver / Sah73 | DPAS-Graph | 0.3405 | 0.3278 | 1.5171 |
| liver / Sah80 | DPAS-Graph | 0.3184 | 0.3045 | 1.0717 |
| liver / Sah73 | HIPI | 0.0297 | 0.0368 | 1.4995 |
| liver / Sah80 | HIPI | 0.0372 | 0.0411 | 1.2170 |
| liver / Sah73 | SPRINT | 0.4290 | 0.3858 | 1.3369 |
| liver / Sah80 | SPRINT | 0.3254 | 0.3378 | 1.0355 |

**Table S2:** All HIPI and SPRINT single-seed runs. The main comparison averages all four seed predictions before computing metrics. Both methods are adapted to the common spot-level protocol.

| Tissue | Method | Seed | SCC | PCC | RMSE |
|---|---|---|---|---|---|
| tonsil | SPRINT | 42 | 0.6177 | 0.6354 | 0.8661 |
| tonsil | SPRINT | 99 | 0.6240 | 0.6369 | 0.8540 |
| tonsil | SPRINT | 7 | 0.6001 | 0.6217 | 0.8480 |
| tonsil | SPRINT | 1024 | 0.6276 | 0.6437 | 0.8352 |
| spleen | SPRINT | 42 | 0.5156 | 0.6076 | 1.2204 |
| spleen | SPRINT | 99 | 0.5151 | 0.5867 | 1.2970 |

Table S2 (continued)

| Tissue | Method | Seed | SCC | PCC | RMSE |
|---|---|---|---|---|---|
| spleen | SPRINT | 7 | 0.5163 | 0.5915 | 1.2580 |
| spleen | SPRINT | 1024 | 0.5029 | 0.5812 | 1.2762 |
| liver | SPRINT | 42 | 0.3156 | 0.3032 | 1.2315 |
| liver | SPRINT | 99 | 0.3421 | 0.3288 | 1.2395 |
| liver | SPRINT | 7 | 0.3328 | 0.3195 | 1.2208 |
| liver | SPRINT | 1024 | 0.3261 | 0.3132 | 1.2272 |
| tonsil | HIPI | 42 | 0.1821 | 0.2272 | 1.1419 |
| tonsil | HIPI | 99 | 0.2206 | 0.2604 | 1.0617 |
| tonsil | HIPI | 7 | 0.2635 | 0.2887 | 1.0212 |
| tonsil | HIPI | 1024 | 0.2477 | 0.2826 | 1.0124 |
| spleen | HIPI | 42 | 0.1871 | 0.1782 | 1.4827 |
| spleen | HIPI | 99 | 0.2027 | 0.1922 | 1.5185 |
| spleen | HIPI | 7 | 0.1855 | 0.1768 | 1.4906 |
| spleen | HIPI | 1024 | 0.1952 | 0.1887 | 1.5032 |
| liver | HIPI | 42 | 0.0206 | 0.0190 | 1.3987 |
| liver | HIPI | 99 | -0.0038 | -0.0028 | 1.3728 |
| liver | HIPI | 7 | 0.0281 | 0.0292 | 1.3859 |
| liver | HIPI | 1024 | -0.0057 | 0.0041 | 1.3782 |

**Table S3:** Matched modality, graph-branch, and fusion controls. Graph removals are separately retrained models. In liver, all seven configurations use the same training recipe.

| Tissue | Configuration | SCC | PCC | RMSE |
|---|---|---|---|---|
| tonsil | RNA-only NN | 0.6136 | 0.6362 | 0.8077 |
| tonsil | RNA+graph | 0.6422 | 0.6593 | 0.8098 |
| tonsil | RNA+image | 0.6356 | 0.6607 | 0.8666 |
| tonsil | No physical graph | 0.6420 | 0.6661 | 0.8647 |
| tonsil | No transcriptomic graph | 0.6490 | 0.6720 | 0.8228 |
| tonsil | Token concatenation | 0.6501 | 0.6727 | 0.8788 |
| tonsil | MorphoProt | 0.6515 | 0.6740 | 0.8215 |
| spleen | RNA-only NN | 0.5638 | 0.6382 | 1.2350 |
| spleen | RNA+graph | 0.5839 | 0.6530 | 1.2174 |
| spleen | RNA+image | 0.5860 | 0.6541 | 1.2234 |
| spleen | No physical graph | 0.5899 | 0.6603 | 1.2300 |
| spleen | No transcriptomic graph | 0.6042 | 0.6699 | 1.1932 |
| spleen | Token concatenation | 0.5940 | 0.6545 | 1.3005 |
| spleen | MorphoProt | 0.6041 | 0.6700 | 1.2345 |
| liver | RNA-only NN | 0.3064 | 0.3283 | 1.2300 |
| liver | RNA+graph | 0.3308 | 0.3453 | 1.1887 |
| liver | RNA+image | 0.3569 | 0.3780 | 1.2168 |
| liver | No physical graph | 0.3600 | 0.3805 | 1.1966 |
| liver | No transcriptomic graph | 0.3674 | 0.3808 | 1.1773 |
| liver | Token concatenation | 0.3688 | 0.3820 | 1.1738 |
| liver | MorphoProt | 0.3717 | 0.3855 | 1.1653 |

**Table S4:** Complete protein panels: MorphoProt versus matched RNA-only.

| Tissue | Protein | MorphoProt | RNA-only | ΔSCC |
|---|---|---|---|---|
| liver | ACTA2 | 0.5904 | 0.5947 | -0.0043 |
| liver | BCL2 | 0.6017 | 0.5327 | 0.0690 |
| liver | CCR7 | 0.3695 | 0.3553 | 0.0142 |
| liver | CD14 | 0.3646 | 0.3198 | 0.0448 |
| liver | CD163 | 0.3851 | 0.3009 | 0.0843 |
| liver | CD19 | 0.2678 | 0.2558 | 0.0120 |
| liver | CD27 | 0.3221 | 0.2858 | 0.0363 |
| liver | CD274 | 0.3902 | 0.1848 | 0.2054 |
| liver | CD3E | 0.4229 | 0.4493 | -0.0264 |
| liver | CD4 | 0.3085 | 0.2866 | 0.0219 |
| liver | CD40 | 0.2456 | 0.1775 | 0.0681 |
| liver | CD68 | 0.5300 | 0.4740 | 0.0560 |
| liver | CD8A | 0.5017 | 0.4397 | 0.0620 |
| liver | CEACAM8 | 0.2216 | 0.0705 | 0.1511 |
| liver | CR2 | 0.2064 | 0.0961 | 0.1102 |
| liver | CXCR5 | 0.2529 | 0.2222 | 0.0307 |
| liver | EPCAM | 0.6347 | 0.6183 | 0.0164 |
| liver | FCGR3A | 0.3570 | 0.2092 | 0.1477 |
| liver | HLA-DRA | 0.1907 | 0.1165 | 0.0742 |
| liver | ITGAM | 0.1168 | -0.1031 | 0.2199 |
| liver | ITGAX | 0.4814 | 0.3721 | 0.1093 |
| liver | KRT5 | 0.7107 | 0.6531 | 0.0576 |
| liver | MS4A1 | 0.2035 | 0.1534 | 0.0501 |
| liver | PAX5 | 0.1267 | 0.1685 | -0.0418 |
| liver | PCNA | 0.3526 | 0.2577 | 0.0949 |
| liver | PDCD1 | 0.2285 | 0.1245 | 0.1039 |
| liver | PECAM1 | 0.4117 | 0.2894 | 0.1223 |
| liver | PTPRC_1 | 0.2417 | 0.1967 | 0.0450 |
| liver | PTPRC_2 | 0.3787 | 0.3048 | 0.0739 |
| liver | SDC1 | 0.3389 | 0.3608 | -0.0219 |
| liver | VIM | 0.7674 | 0.7299 | 0.0375 |
| spleen | B220_CD45R | 0.7702 | 0.7044 | 0.0657 |
| spleen | CD105 | 0.5837 | 0.5578 | 0.0259 |
| spleen | CD11b | 0.5564 | 0.5178 | 0.0385 |
| spleen | CD163 | 0.7471 | 0.7015 | 0.0456 |
| spleen | CD169_siglec | 0.6770 | 0.5931 | 0.0839 |
| spleen | CD19 | 0.7824 | 0.7081 | 0.0743 |
| spleen | CD20 | 0.3772 | 0.3815 | -0.0044 |
| spleen | CD29 | 0.6981 | 0.6496 | 0.0486 |
| spleen | CD3 | 0.6773 | 0.6130 | 0.0643 |
| spleen | CD31 | 0.5751 | 0.5482 | 0.0269 |
| spleen | CD38 | 0.6020 | 0.5594 | 0.0426 |
| spleen | CD4 | 0.6970 | 0.6473 | 0.0497 |
| spleen | CD68 | 0.7427 | 0.6842 | 0.0585 |
| spleen | CD8 | 0.6264 | 0.5947 | 0.0317 |
| spleen | EpCAM | 0.3204 | 0.3155 | 0.0049 |

Table S4 (continued)

| Tissue | Protein | MorphoProt | RNA-only | ΔSCC |
|---|---|---|---|---|
| spleen | F4_80 | 0.8064 | 0.7472 | 0.0591 |
| spleen | IgD | 0.7841 | 0.7145 | 0.0696 |
| spleen | IgM | 0.4734 | 0.4255 | 0.0479 |
| spleen | Ly6C | 0.3925 | 0.3855 | 0.0070 |
| spleen | Ly6G | 0.4089 | 0.4026 | 0.0063 |
| spleen | MadCAM1 | 0.3880 | 0.3892 | -0.0012 |
| tonsil | ACTA2-1 | 0.8413 | 0.8081 | 0.0332 |
| tonsil | BCL2-1 | 0.6149 | 0.6253 | -0.0104 |
| tonsil | CCR7-1 | 0.4828 | 0.4906 | -0.0077 |
| tonsil | CD14-1 | 0.1547 | 0.2343 | -0.0796 |
| tonsil | CD163-1 | 0.4789 | 0.4405 | 0.0384 |
| tonsil | CD19-1 | 0.7664 | 0.7509 | 0.0155 |
| tonsil | CD27-1 | 0.4178 | 0.3894 | 0.0284 |
| tonsil | CD274-1 | 0.7639 | 0.7324 | 0.0315 |
| tonsil | CD3E-1 | 0.9291 | 0.9266 | 0.0024 |
| tonsil | CD4-1 | 0.6724 | 0.6551 | 0.0172 |
| tonsil | CD40-1 | 0.2219 | 0.0573 | 0.1647 |
| tonsil | CD68-1 | 0.6200 | 0.5542 | 0.0659 |
| tonsil | CD8A-1 | 0.8841 | 0.8710 | 0.0131 |
| tonsil | CEACAM8-1 | 0.5678 | 0.5306 | 0.0372 |
| tonsil | CR2-1 | 0.5119 | 0.5170 | -0.0051 |
| tonsil | CXCR5-1 | 0.9147 | 0.9012 | 0.0135 |
| tonsil | EPCAM-1 | 0.4809 | 0.4365 | 0.0444 |
| tonsil | FCGR3A-1 | 0.6260 | 0.6138 | 0.0122 |
| tonsil | HLA-DRA | 0.7213 | 0.7041 | 0.0172 |
| tonsil | ITGAM-1 | 0.5172 | 0.5416 | -0.0245 |
| tonsil | ITGAX-1 | 0.7476 | 0.6679 | 0.0797 |
| tonsil | KRT5-1 | 0.8428 | 0.8249 | 0.0178 |
| tonsil | MS4A1-1 | 0.8220 | 0.6740 | 0.1479 |
| tonsil | PAX5-1 | 0.8622 | 0.8593 | 0.0029 |
| tonsil | PCNA-1 | 0.6225 | 0.6029 | 0.0195 |
| tonsil | PDCD1-1 | 0.8296 | 0.8101 | 0.0195 |
| tonsil | PECAM1-1 | 0.8404 | 0.7810 | 0.0595 |
| tonsil | SDC1-1 | 0.4384 | 0.1846 | 0.2538 |
| tonsil | VIM-1 | 0.7013 | 0.6097 | 0.0916 |

**Table S5:** Spot-level matched cognate-RNA panels: 22 tonsil and 21 spleen proteins. B220/CD45R uses total Ptprc expression; it is not an isoform-specific measurement.

| Tissue | Protein | MorphoProt | RNA-only | RNA proxy |
|---|---|---|---|---|
| spleen | B220_CD45R | 0.7702 | 0.7044 | 0.3794 |
| spleen | CD105 | 0.5837 | 0.5578 | 0.0452 |
| spleen | CD11b | 0.5564 | 0.5178 | 0.0311 |
| spleen | CD163 | 0.7471 | 0.7015 | 0.0493 |
| spleen | CD169_siglec | 0.6770 | 0.5931 | 0.2187 |

Table S5 (continued)

| Tissue | Protein | MorphoProt | RNA-only | RNA proxy |
|---|---|---|---|---|
| spleen | CD19 | 0.7824 | 0.7081 | 0.4751 |
| spleen | CD20 | 0.3772 | 0.3815 | 0.1993 |
| spleen | CD29 | 0.6981 | 0.6496 | 0.0372 |
| spleen | CD3 | 0.6773 | 0.6130 | 0.2916 |
| spleen | CD31 | 0.5751 | 0.5482 | 0.2395 |
| spleen | CD38 | 0.6020 | 0.5594 | 0.2709 |
| spleen | CD4 | 0.6970 | 0.6473 | 0.2942 |
| spleen | CD68 | 0.7427 | 0.6842 | 0.0164 |
| spleen | CD8 | 0.6264 | 0.5947 | 0.2613 |
| spleen | EpCAM | 0.3204 | 0.3155 | 0.0243 |
| spleen | F4_80 | 0.8064 | 0.7472 | 0.0586 |
| spleen | IgD | 0.7841 | 0.7145 | 0.5431 |
| spleen | IgM | 0.4734 | 0.4255 | 0.3450 |
| spleen | Ly6C | 0.3925 | 0.3855 | 0.0992 |
| spleen | Ly6G | 0.4089 | 0.4026 | 0.0294 |
| spleen | MadCAM1 | 0.3880 | 0.3892 | 0.1996 |
| tonsil | ACTA2-1 | 0.8413 | 0.8081 | 0.3104 |
| tonsil | BCL2-1 | 0.6149 | 0.6253 | 0.3347 |
| tonsil | CCR7-1 | 0.4828 | 0.4906 | 0.3449 |
| tonsil | CD14-1 | 0.1547 | 0.2343 | 0.1102 |
| tonsil | CD163-1 | 0.4789 | 0.4405 | 0.3597 |
| tonsil | CD19-1 | 0.7664 | 0.7509 | 0.6520 |
| tonsil | CD274-1 | 0.7639 | 0.7324 | 0.2221 |
| tonsil | CD3E-1 | 0.9291 | 0.9266 | 0.7358 |
| tonsil | CD4-1 | 0.6724 | 0.6551 | 0.4708 |
| tonsil | CD40-1 | 0.2219 | 0.0573 | 0.0733 |
| tonsil | CD8A-1 | 0.8841 | 0.8710 | 0.4756 |
| tonsil | CR2-1 | 0.5119 | 0.5170 | 0.4606 |
| tonsil | CXCR5-1 | 0.9147 | 0.9012 | 0.5803 |
| tonsil | EPCAM-1 | 0.4809 | 0.4365 | 0.3239 |
| tonsil | FCGR3A-1 | 0.6260 | 0.6138 | 0.2112 |
| tonsil | ITGAX-1 | 0.7476 | 0.6679 | 0.2844 |
| tonsil | KRT5-1 | 0.8428 | 0.8249 | 0.7406 |
| tonsil | PAX5-1 | 0.8622 | 0.8593 | 0.7108 |
| tonsil | PCNA-1 | 0.6225 | 0.6029 | 0.2636 |
| tonsil | PDCD1-1 | 0.8296 | 0.8101 | 0.4375 |
| tonsil | PECAM1-1 | 0.8404 | 0.7810 | 0.7041 |
| tonsil | SDC1-1 | 0.4384 | 0.1846 | 0.4600 |

**Table S6:** Liver section results for all retained methods and controls. Pooled liver SCC is recomputed across both sections and is not the average of these values.

| Section | Method / configuration | SCC | PCC | RMSE |
|---|---|---|---|---|
| Sah73 | DGAT | 0.3165 | 0.3106 | 1.4001 |
| Sah80 | DGAT | 0.2552 | 0.2562 | 1.0961 |

Table S6 (continued)

| Section | Method / configuration | SCC | PCC | RMSE |
|---|---|---|---|---|
| Sah73 | DPAS-Graph | 0.3405 | 0.3278 | 1.5171 |
| Sah80 | DPAS-Graph | 0.3184 | 0.3045 | 1.0717 |
| Sah73 | RNA+image | 0.4171 | 0.4166 | 1.3601 |
| Sah80 | RNA+image | 0.3469 | 0.3579 | 1.0227 |
| Sah73 | RNA+graph | 0.3204 | 0.3210 | 1.3415 |
| Sah80 | RNA+graph | 0.3536 | 0.3670 | 0.9788 |
| Sah73 | MorphoProt | 0.3918 | 0.3898 | 1.3086 |
| Sah80 | MorphoProt | 0.3660 | 0.3853 | 0.9698 |
| Sah73 | RNA-only NN | 0.3236 | 0.3292 | 1.3732 |
| Sah80 | RNA-only NN | 0.3322 | 0.3316 | 1.0363 |
| Sah73 | No physical graph | 0.4192 | 0.4183 | 1.3382 |
| Sah80 | No physical graph | 0.3386 | 0.3527 | 1.0045 |
| Sah73 | No transcriptomic graph | 0.3854 | 0.3846 | 1.3267 |
| Sah80 | No transcriptomic graph | 0.3636 | 0.3851 | 0.9725 |
| Sah73 | Token concatenation | 0.4044 | 0.4044 | 1.3103 |
| Sah80 | Token concatenation | 0.3686 | 0.3878 | 0.9894 |
| Sah73 | HIPI | 0.0297 | 0.0368 | 1.4995 |
| Sah80 | HIPI | 0.0372 | 0.0411 | 1.2170 |
| Sah73 | SPRINT | 0.4290 | 0.3858 | 1.3369 |
| Sah80 | SPRINT | 0.3254 | 0.3378 | 1.0355 |

**Table S7:** Validation-selected ridge-regression controls. Their objectives and function classes differ from the neural controls.

| Tissue | Control | SCC | PCC | RMSE |
|---|---|---|---|---|
| tonsil | RNA ridge | 0.6063 | 0.6196 | 0.8327 |
| tonsil | RNA-PCA ridge | 0.5984 | 0.6089 | 0.8822 |
| tonsil | PCA+spatial ridge | 0.5834 | 0.5879 | 0.8892 |
| spleen | RNA ridge | 0.5241 | 0.5497 | 1.3344 |
| spleen | RNA-PCA ridge | 0.5393 | 0.5376 | 1.3444 |
| spleen | PCA+spatial ridge | 0.5613 | 0.5589 | 1.3254 |
| liver | RNA ridge | 0.3088 | 0.2950 | 1.2270 |
| liver | RNA-PCA ridge | 0.3235 | 0.3013 | 1.2194 |
| liver | PCA+spatial ridge | 0.2961 | 0.2797 | 1.2352 |

**Table S8:** Fusion comparison and trainable parameter counts. Frozen image feature extractors are excluded. Token concatenation adds 1,317,887 parameters in every tissue.

| Tissue | Cross SCC | Concat SCC | Cross params | Concat params |
|---|---|---|---|---|
| tonsil | 0.6515 | 0.6501 | 6,718,610 | 8,036,497 |
| spleen | 0.6041 | 0.5940 | 5,405,426 | 6,723,313 |
| liver | 0.3717 | 0.3688 | 5,408,526 | 6,726,413 |

**Table S9:** Single-field models. The full model is retrained with one field of view (four seeds). Validation SCC is the mean over seeds; test SCC is that of the four-seed ensemble. Δ is five-field minus single-field SCC with a paired bootstrap 95% CI over proteins; 'improved' counts proteins better predicted by the five-field model; 'retained' is the fraction of the histology gain (five-field minus no-image SCC) reached by the single field. Field sizes in $\mu$m use 0.215 (tonsil), 4.82 (spleen) and 4.19 (liver, mean of test sections) $\mu$m per pixel. Five-field SCC: 0.6515, 0.6041 and 0.3717. A per-protein oracle that takes, for each protein, the highest test SCC over the five single-field ensembles reaches 0.600 in spleen and 0.369 in liver, below the five-field model, so the gain comes from combining fields rather than from selecting one; any single field retains 21–57% (spleen) and 45–74% (liver) of the histology gain.

| Tissue | px | $\mu$m | Val SCC | Test SCC | Δ [95% CI] | Improved | Retained |
|---|---|---|---|---|---|---|---|
| tonsil | 256 | 55 | 0.8473 | 0.6492 | +0.0024 [-0.0030, +0.0081] | 15/29 | 0.75 |
| tonsil | 512 | 110 | 0.8524 | 0.6483 | +0.0032 [-0.0025, +0.0103] | 14/29 | 0.66 |
| tonsil | 768 | 165 | 0.8523 | 0.6545 | -0.0030 [-0.0091, +0.0026] | 11/29 | 1.32 |
| tonsil | 1024 | 220 | 0.8527 | 0.6539 | -0.0024 [-0.0099, +0.0046] | 14/29 | 1.25 |
| tonsil | 1280 | 275 | 0.8533 | 0.6509 | +0.0007 [-0.0078, +0.0089] | 16/29 | 0.93 |
| spleen | 16 | 77 | 0.5941 | 0.5899 | +0.0142 [+0.0070, +0.0209] | 18/21 | 0.30 |
| spleen | 32 | 154 | 0.5999 | 0.5923 | +0.0118 [+0.0056, +0.0175] | 17/21 | 0.41 |
| spleen | 64 | 308 | 0.6021 | 0.5954 | +0.0087 [+0.0050, +0.0124] | 17/21 | 0.57 |
| spleen | 96 | 463 | 0.6052 | 0.5888 | +0.0153 [+0.0114, +0.0191] | 19/21 | 0.24 |
| spleen | 128 | 617 | 0.6053 | 0.5881 | +0.0160 [+0.0122, +0.0198] | 20/21 | 0.21 |
| liver | 16 | 67 | 0.6646 | 0.3510 | +0.0207 [+0.0128, +0.0285] | 26/31 | 0.49 |
| liver | 32 | 134 | 0.6646 | 0.3562 | +0.0155 [+0.0077, +0.0230] | 27/31 | 0.62 |
| liver | 64 | 268 | 0.6685 | 0.3493 | +0.0223 [+0.0130, +0.0315] | 26/31 | 0.45 |
| liver | 96 | 402 | 0.6697 | 0.3585 | +0.0132 [+0.0057, +0.0207] | 23/31 | 0.68 |
| liver | 128 | 536 | 0.6746 | 0.3610 | +0.0107 [+0.0041, +0.0170] | 23/31 | 0.74 |

**Table S10:** Image-encoder control. Tonsil is retrained with ImageNet ResNet18 features and liver with UNI2-H features, keeping everything else fixed; the paper encoders are shown for reference. Δ is five-field minus single-field SCC (paired 95% CI over proteins). No-image SCC: tonsil 0.6422, liver 0.3308.

| Tissue | Encoder | Fields | Test SCC | Δ [95% CI] | Improved |
|---|---|---|---|---|---|
| tonsil | UNI2-H (paper) | five fields | 0.6515 | – | – |
| | | 256 px | 0.6492 | +0.0024 [-0.0030, +0.0081] | 15/29 |
| | | 512 px | 0.6483 | +0.0032 [-0.0025, +0.0103] | 14/29 |
| | | 768 px | 0.6545 | -0.0030 [-0.0091, +0.0026] | 11/29 |
| | | 1024 px | 0.6539 | -0.0024 [-0.0099, +0.0046] | 14/29 |
| | | 1280 px (val.) | 0.6509 | +0.0007 [-0.0078, +0.0089] | 16/29 |
| tonsil | ResNet18 | five fields | 0.6418 | – | – |
| | | 256 px | 0.6384 | +0.0034 [-0.0079, +0.0176] | 14/29 |
| | | 512 px (val.) | 0.6429 | -0.0011 [-0.0090, +0.0078] | 11/29 |
| | | 768 px | 0.6412 | +0.0006 [-0.0079, +0.0110] | 13/29 |
| | | 1024 px | 0.6441 | -0.0023 [-0.0123, +0.0097] | 9/29 |
| | | 1280 px | 0.6484 | -0.0066 [-0.0158, +0.0038] | 9/29 |
| liver | ResNet18 (paper) | five fields | 0.3717 | – | – |
| | | 16 px | 0.3510 | +0.0207 [+0.0128, +0.0285] | 26/31 |
| | | 32 px | 0.3562 | +0.0155 [+0.0077, +0.0230] | 27/31 |
| | | 64 px | 0.3493 | +0.0223 [+0.0130, +0.0315] | 26/31 |
| | | 96 px | 0.3585 | +0.0132 [+0.0057, +0.0207] | 23/31 |

Table S10 (continued)

| Tissue | Encoder | Fields | Test SCC | Δ [95% CI] | Improved |
|---|---|---|---|---|---|
| | | 128 px (val.) | 0.3610 | +0.0107 [+0.0041, +0.0170] | 23/31 |
| liver | UNI2-H | five fields | 0.3256 | – | – |
| | | 16 px | 0.3665 | -0.0409 [-0.0837, -0.0027] | 12/31 |
| | | 32 px | 0.3681 | -0.0425 [-0.0716, -0.0152] | 8/31 |
| | | 64 px | 0.3544 | -0.0288 [-0.0455, -0.0132] | 6/31 |
| | | 96 px | 0.3471 | -0.0215 [-0.0352, -0.0087] | 8/31 |
| | | 128 px (val.) | 0.3248 | +0.0008 [-0.0080, +0.0098] | 17/31 |

**Table S11:** Field-of-view sensitivity of the full five-field model (four-seed ensembles; same seeds, splits and training recipe as the corresponding paper model). Physically matched fields use the same micrometer sizes in both tissues; in liver the two smallest fields span 10–28 pixels of the high-resolution image. Mean pairwise CKA between the five field embeddings of the test spots: tonsil UNI2-H 0.80, tonsil wide fields 0.55, tonsil ResNet18 0.56 (female) and 0.68 (male section), spleen ResNet18 0.21, liver ResNet18 0.40.

| Tissue | Fields | SCC | Δ vs paper |
|---|---|---|---|
| tonsil | paper fields (55–275 $\mu$m) | 0.6515 | – |
| tonsil | wide fields (21–434 $\mu$m) | 0.6488 | −0.0027 |
| tonsil | physical fields (50, 100, 150, 200, 600 $\mu$m) | 0.6459 | -0.0056 |
| liver | paper fields (67–536 $\mu$m) | 0.3717 | – |
| liver | physical fields (50, 100, 150, 200, 600 $\mu$m) | 0.3513 | -0.0204 |
| liver Sah73 | physical fields | 0.3563 | -0.0355 |
| liver Sah80 | physical fields | 0.3685 | +0.0025 |

**Table S12:** Protein-specific attention queries. Each variant replaces or augments the shared spot-level query by one learned query per protein; Δ is relative to the paper model (shared query; tonsil 0.6515, spleen 0.6041) with a paired 95% CI over proteins; 'improved' counts proteins better predicted by the variant.

| Tissue | Variant | SCC | Δ [95% CI] | Improved |
|---|---|---|---|---|
| tonsil | protein queries added to the shared query | 0.6460 | -0.0056 [-0.0109, -0.0009] | 9/29 |
| tonsil | protein queries, zero-initialized residual | 0.6490 | -0.0025 [-0.0079, +0.0031] | 11/29 |
| tonsil | protein queries only | 0.6446 | -0.0069 [-0.0246, +0.0105] | 7/29 |
| spleen | protein queries added to the shared query | 0.5968 | -0.0073 [-0.0127, -0.0017] | 5/21 |
| spleen | protein queries, zero-initialized residual | 0.5980 | -0.0061 [-0.0098, -0.0024] | 6/21 |
| spleen | protein queries only | 0.5789 | -0.0252 [-0.0354, -0.0153] | 3/21 |

**Table S13:** Spleen: Spearman correlation between the per-spot scale preference (attention on the two finest minus the two coarsest fields, head mean) and each measured protein, for each of the four seeds. Attention entropy normalized by $\log 5$: spleen 0.88, tonsil 0.96; seed-to-seed correlation of the per-spot preference: spleen 0.52, tonsil 0.07.

| Protein | Mean $\rho$ | Min | Max | Same sign |
|---|---|---|---|---|
| CD4 | +0.554 | +0.527 | +0.585 | yes |
| IgD | +0.498 | +0.457 | +0.545 | yes |

Table S13 (continued)

| Protein | Mean $\rho$ | Min | Max | Same sign |
|---|---|---|---|---|
| B220 | +0.497 | +0.463 | +0.540 | yes |
| CD19 | +0.490 | +0.445 | +0.537 | yes |
| CD3 | +0.488 | +0.459 | +0.531 | yes |
| CD169 | +0.350 | +0.329 | +0.383 | yes |
| CD8 | +0.308 | +0.275 | +0.373 | yes |
| CD38 | +0.244 | +0.203 | +0.267 | yes |
| MadCAM1 | +0.232 | +0.210 | +0.245 | yes |
| IgM | +0.174 | +0.152 | +0.186 | yes |
| Ly6G | +0.166 | +0.144 | +0.200 | yes |
| Ly6C | +0.155 | +0.134 | +0.178 | yes |
| CD31 | +0.150 | +0.130 | +0.199 | yes |
| CD20 | +0.128 | +0.114 | +0.167 | yes |
| EpCAM | +0.097 | +0.084 | +0.122 | yes |
| CD11b | +0.063 | +0.040 | +0.117 | yes |
| CD105 | -0.019 | -0.042 | +0.036 | no |
| CD29 | -0.179 | -0.217 | -0.103 | yes |
| CD68 | -0.268 | -0.312 | -0.184 | yes |
| CD163 | -0.399 | -0.420 | -0.343 | yes |
| F4/80 | -0.447 | -0.477 | -0.375 | yes |

**Table S14:** Downstream analyses of the multiscale and fusion controls, using the protocol of Fig. 4 (matched panels, $k$-means over ten seeds). Mean ARI is averaged over $k = 3, \ldots, 8$.

| Tissue | Representation | ARI ($k = 5$) | ARI (mean $k$) | Moran's I $r$ |
|---|---|---|---|---|
| tonsil | MorphoProt | 0.703 | 0.652 | 0.817 |
| tonsil | Best single field | 0.700 | 0.654 | 0.837 |
| tonsil | Token concat | 0.688 | 0.642 | 0.806 |
| tonsil | No image | 0.677 | 0.647 | 0.768 |
| tonsil | SPRINT | 0.513 | 0.607 | 0.825 |
| tonsil | RNA-only NN | 0.656 | 0.620 | 0.776 |
| spleen | MorphoProt | 0.334 | 0.256 | 0.915 |
| spleen | Best single field | 0.339 | 0.262 | 0.924 |
| spleen | Token concat | 0.258 | 0.243 | 0.926 |
| spleen | No image | 0.330 | 0.247 | 0.897 |
| spleen | SPRINT | 0.195 | 0.170 | 0.780 |
| spleen | RNA-only NN | 0.280 | 0.212 | 0.849 |

**Table S15:** Germinal-center identification by the cluster-free eight-marker score: AUPRC with 95% spatial-block bootstrap CIs (74 blocks of eight spot pitches, about 0.8 mm; 2,000 resamples). GC prevalence 0.080. The mRNA score requires the RNA matrix and has no CI here.

| Representation | AUPRC | 95% CI |
|---|---|---|
| Measured protein | 0.702 | [0.621, 0.774] |

Table S15 (continued)

| Representation | AUPRC | 95% CI |
|---|---|---|
| MorphoProt | 0.786 | [0.701, 0.854] |
| SPRINT | 0.800 | [0.725, 0.861] |
| DGAT | 0.771 | [0.718, 0.820] |
| RNA-only | 0.755 | [0.692, 0.813] |
| DPAS-Graph | 0.744 | [0.692, 0.798] |
| HIPI | 0.639 | [0.548, 0.715] |
| Cognate RNA | 0.578 | [0.486, 0.666] |
| mRNA (7/8 marker genes) | 0.398 | – |

**Table S16: Recovery of pathologist-annotated germinal centers on the held-out tonsil section.** Cluster-based metrics are means over ten seeds at $k = 5$; the cluster with the highest F1 against the annotation is called GC. Sig. AUROC/AUPRC use the cluster-free eight-marker GC signature (identical in both blocks). Bold: best among predictors and transcript-level representations (measured proteins excluded). GC prevalence: 391/4,906 spots (0.080).

| Representation | F1 | Prec. | Recall | MCC | Jaccard | $\kappa$ | Bal. acc. | Sig. AUROC | Sig. AUPRC |
|---|---|---|---|---|---|---|---|---|---|
| *k-means, $k = 5$* | | | | | | | | | |
| Measured protein | 0.711 | 0.674 | 0.752 | 0.686 | 0.552 | 0.685 | 0.860 | 0.953 | 0.702 |
| MorphoProt | 0.734 | **0.724** | 0.744 | 0.711 | 0.580 | **0.711** | 0.860 | **0.973** | 0.786 |
| SPRINT | **0.735** | 0.712 | 0.760 | **0.712** | **0.581** | **0.711** | 0.867 | 0.972 | **0.800** |
| RNA-only NN | 0.726 | 0.696 | 0.759 | 0.702 | 0.570 | 0.701 | 0.865 | 0.963 | 0.755 |
| DGAT | 0.720 | 0.688 | 0.754 | 0.695 | 0.562 | 0.694 | 0.862 | 0.965 | 0.771 |
| DPAS-Graph | 0.710 | 0.713 | 0.706 | 0.684 | 0.550 | 0.684 | 0.841 | 0.962 | 0.744 |
| HIPI | 0.602 | 0.570 | 0.638 | 0.567 | 0.431 | 0.565 | 0.798 | 0.873 | 0.638 |
| mRNA (3,500 HVG) | 0.524 | 0.362 | **0.949** | 0.536 | 0.355 | 0.462 | **0.902** | 0.919 | 0.398 |
| RNA proxy | 0.466 | 0.309 | 0.944 | 0.480 | 0.303 | 0.393 | 0.880 | 0.940 | 0.578 |
| *Leiden, resolution tuned to $k = 5$ (five seeds)* | | | | | | | | | |
| Measured protein | 0.606 | 0.483 | 0.855 | 0.594 | 0.441 | 0.559 | 0.881 | 0.953 | 0.702 |
| MorphoProt | 0.723 | **0.703** | 0.759 | 0.702 | 0.566 | 0.698 | 0.865 | **0.973** | 0.786 |
| SPRINT | **0.735** | 0.673 | 0.826 | **0.718** | **0.581** | **0.710** | 0.895 | 0.972 | **0.800** |
| RNA-only NN | 0.579 | 0.471 | 0.864 | 0.576 | 0.417 | 0.527 | 0.876 | 0.963 | 0.755 |
| DGAT | 0.717 | 0.652 | 0.797 | 0.694 | 0.559 | 0.690 | 0.880 | 0.965 | 0.771 |
| DPAS-Graph | 0.641 | 0.622 | 0.753 | 0.628 | 0.483 | 0.604 | 0.841 | 0.962 | 0.744 |
| HIPI | 0.584 | 0.546 | 0.660 | 0.553 | 0.416 | 0.542 | 0.801 | 0.873 | 0.638 |
| mRNA (3,500 HVG) | 0.698 | 0.584 | 0.869 | 0.683 | 0.536 | 0.666 | **0.908** | 0.919 | 0.398 |
| RNA proxy | 0.459 | 0.304 | **0.944** | 0.474 | 0.298 | 0.385 | 0.878 | 0.940 | 0.578 |

Accuracy and specificity exceed 0.81 for every representation because 92% of spots are non-GC and are not shown. Balanced accuracy favors representations that over-call GCs (high recall, low precision), so MCC is used as the primary imbalance-robust summary.

### Supplementary Figures

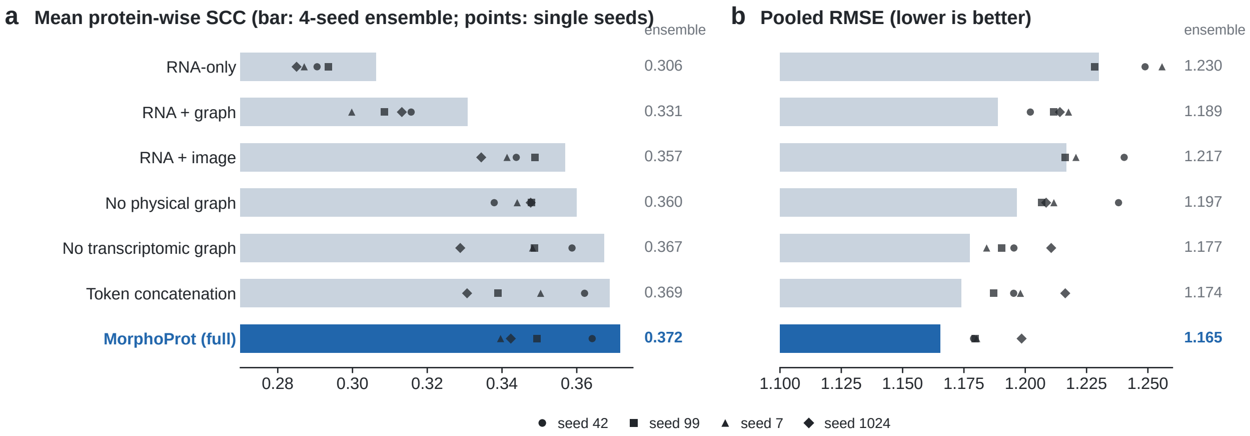


**Figure S1:** Liver matched controls. (a) Mean protein-wise SCC and (b) pooled RMSE. Bars and the numbers on the right are metrics of the four-seed prediction ensemble; markers show the individual training seeds (not biological replicates). All six controls use the same recipe as the full model.

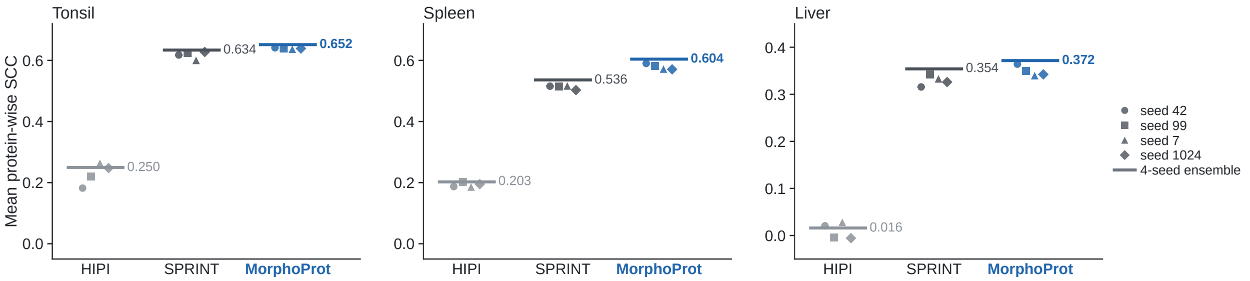


**Figure S2:** Every training seed of HIPI, SPRINT and MorphoProt under the common spot-level protocol. Markers are single-seed SCCs; the horizontal line and number give the SCC of the four-seed prediction ensemble, which is not the mean of the four seed SCCs. No seed is selected for the primary comparison.

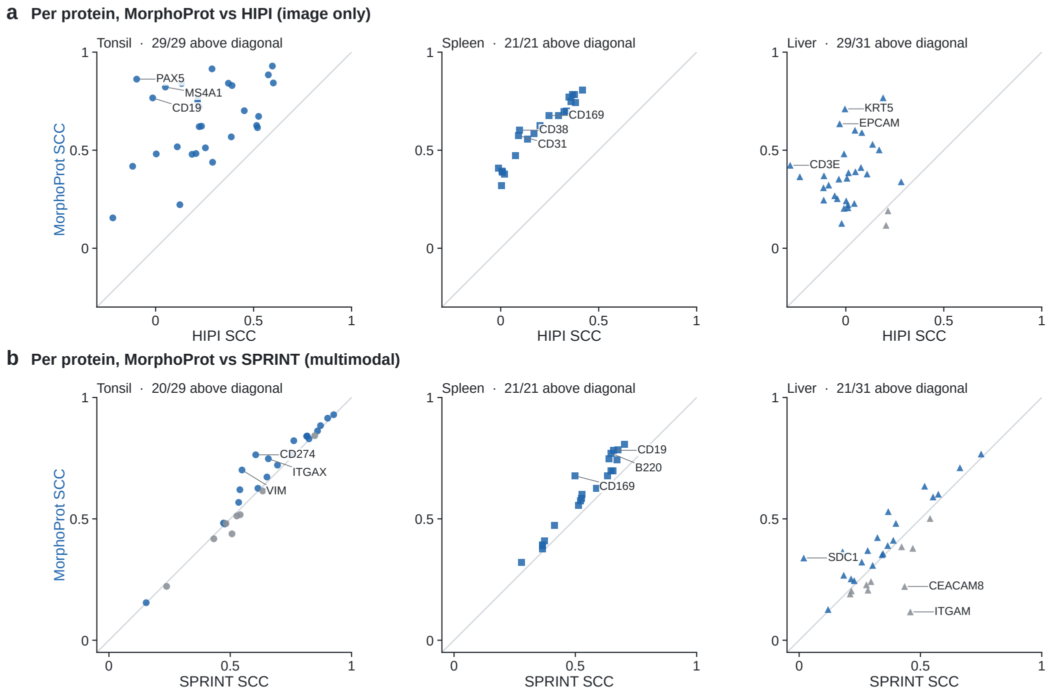


**Figure S3:** Per-protein test SCC of MorphoProt versus (a) image-only HIPI and (b) multimodal SPRINT. Every protein is shown; blue marks proteins above the diagonal (MorphoProt higher), grey those below. The three proteins with the largest absolute difference in each panel are labeled.

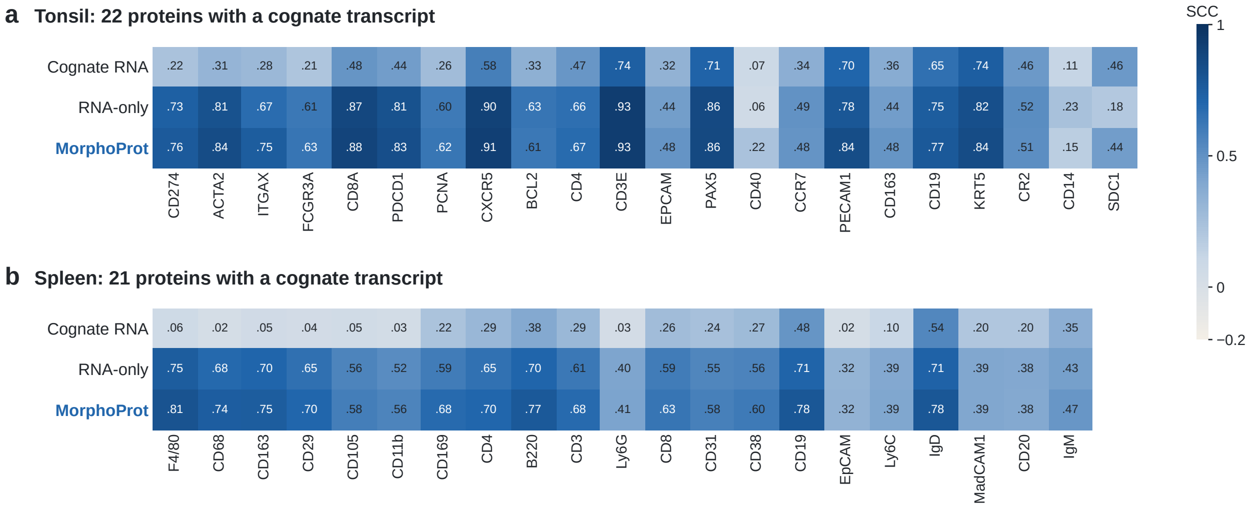


**Figure S4:** Protein-wise SCC of the cognate transcript, the RNA-only network and MorphoProt on the matched panels of (a) tonsil and (b) spleen. Proteins are ordered by MorphoProt minus cognate-RNA SCC; cell values are SCCs on a common color scale from −0.2 to 1. RNA-only is the multivariate neural model trained on the full transcriptome.

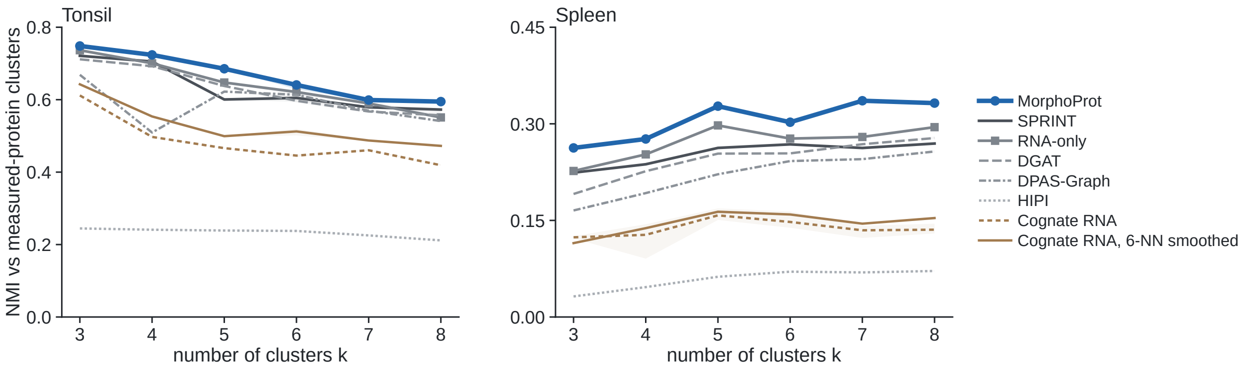


**Figure S5:** Normalized mutual information between $k$-means clusters of each representation and clusters of the measured proteins, on the matched panels, for $k = 3$–8. Lines are means over ten $k$-means seeds; shading spans their minimum and maximum. All values of $k$ are shown.

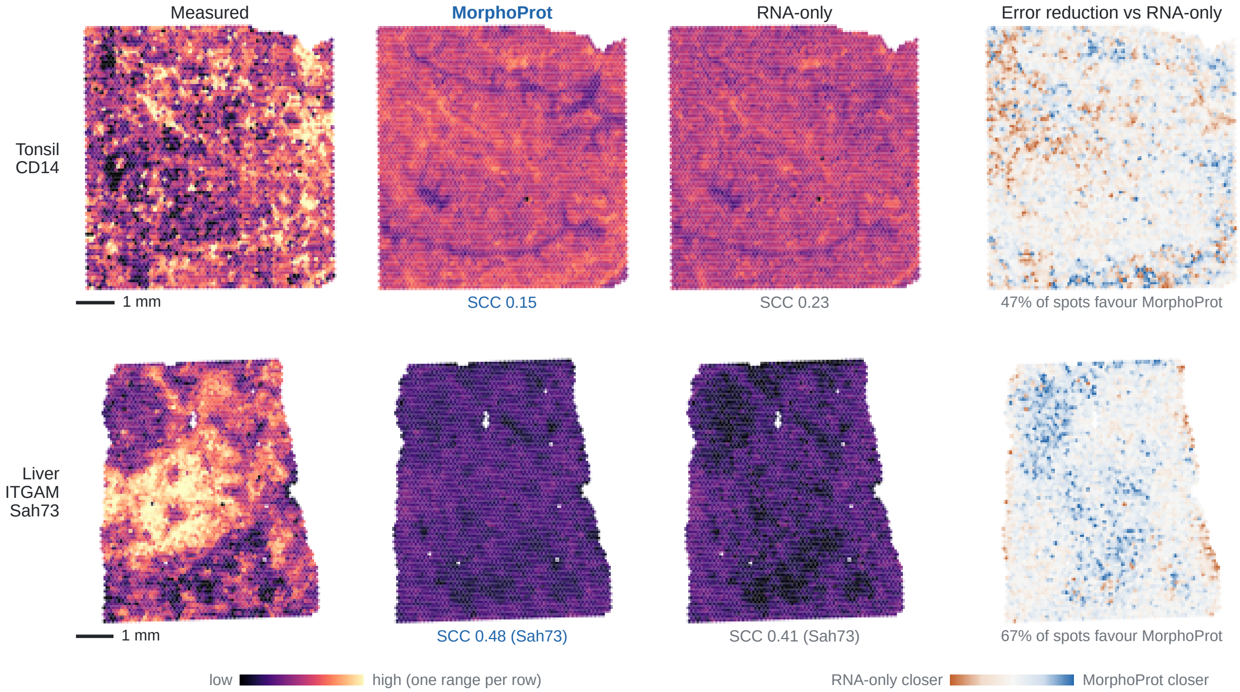


**Figure S6:** Additional spatial cases: tonsil CD14, where RNA-only is closer to the measurement, and liver ITGAM on section Sah73, where MorphoProt is closer. Protein maps in a row share one color range (first–99th percentile of the pooled values). The right column shows absolute RNA-only error minus absolute MorphoProt error per spot (blue: MorphoProt closer), with a symmetric 99th-percentile display limit. SCCs use unclipped values, over all tonsil spots or over Sah73 spots. Scale bar, 1 mm.

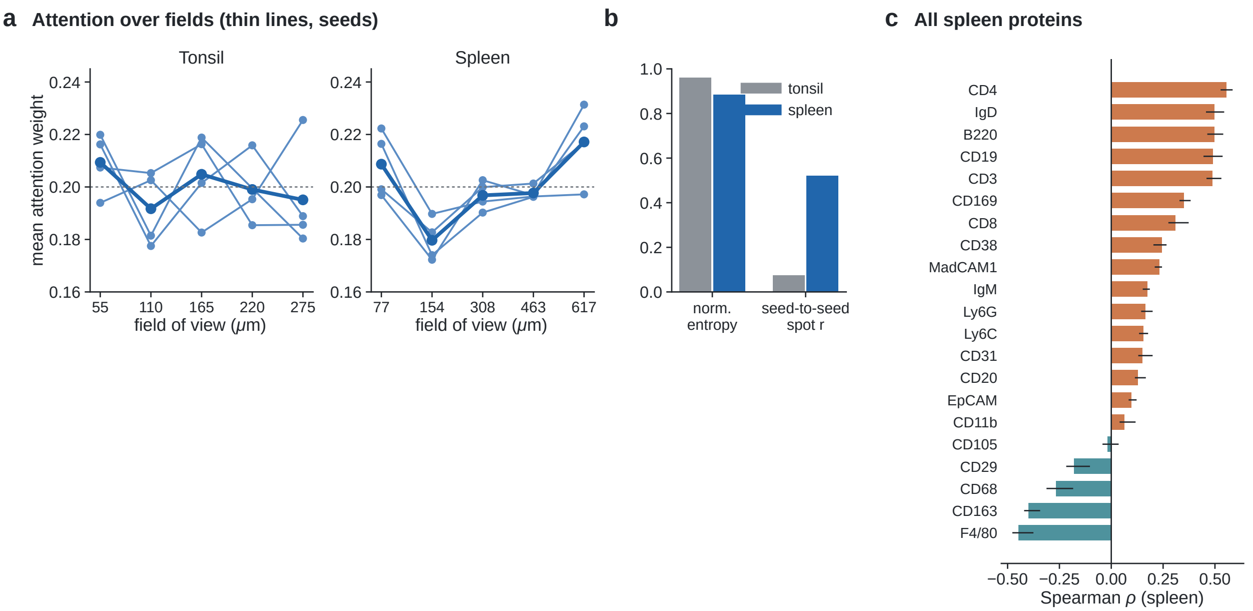


**Figure S7:** Attention over fields of view. **a**, Mean attention weight per field (head mean over test spots) for each of the four seeds (thin lines) and their mean (thick); the dotted line marks uniform weight (0.2). **b**, Normalized attention entropy and seed-to-seed correlation of the per-spot scale preference. **c**, Spearman correlation between spleen scale preference and every measured protein (bars, mean over seeds; lines, range).

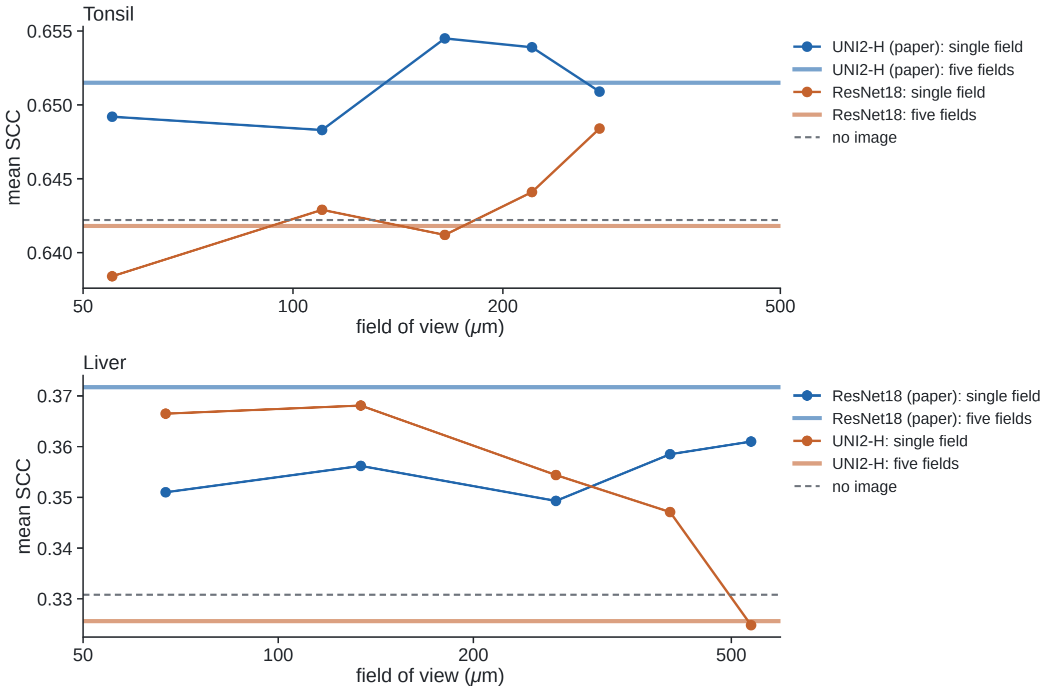


**Figure S8:** Image-encoder control. Single-field (points) and five-field (horizontal lines) test SCC with the paper encoder and with the alternative encoder; the dashed line is the model without histology. Values are listed in Table S10.
